# microRNA-21 promotes dysregulated lipid metabolism and hepatocellular carcinoma

**DOI:** 10.1101/2025.07.19.665686

**Authors:** Chad VanSant-Webb, Jessye C. Castro, Audrey Y. Su, Kiandra Hawkins, Aavrati Saxena, Richard Smith, Jillian Wright, Warren P. Voth, Carrie Barton, Chris Stubben, Ryan M. O’Connell, Gregory S. Ducker, Kimberley J. Evason

**Author notes:** Equal contributions. Co-corresponding: Kimberley J. Evason, 2000 Circle of Hope Way, Research North, Huntsman Cancer Institute, Salt Lake City, UT 84112; Gregory S. Ducker, 15 North Medical Drive East, Salt Lake City, UT 84101. **Conflicts of interest:** Nothing to report. **Statement of Ethics:** Zebrafish (Danio rerio) studies were performed in compliance with the University of Utah Institutional Animal Care and Use Committee guidelines (Protocols #1809 and #2233) under the supervision of Office of Comparative Medicine veterinarians and staff. Studies on human tissues were reviewed and deemed exempt by the University of Utah Institutional Review Board (IRB #00091019).

## Abstract

**Backgrounds and Aims:** The prevalence of hepatocellular carcinoma (HCC) is rising in parallel with increasing obesity and metabolic dysfunction-associated steatohepatitis (MASH). MicroRNAs are key post-transcriptional regulators of gene expression and are attractive targets for HCC therapy. Here we sought to identify and characterize dysregulated microRNAs in MASH-driven HCC (MASH-HCC).

**Approach and Results:** We profiled microRNA expression in liver tissue from patients with MASH and/or MASH-HCC and in zebrafish HCC driven by activated β-catenin (ABC), one of the most commonly mutated oncogenes in MASH-HCC. We found significant overlap between dysregulated human and zebrafish miRNAs, including miR-21, which was increasingly upregulated from normal liver to MASH to MASH-HCC. We generated transgenic zebrafish that overexpress or sponge (downregulate) miR-21. We found that miR-21 overexpression caused larval liver overgrowth and increased HCC while miR-21 sponge suppressed β-catenin-driven larval liver overgrowth. By performing histologic and lipidomic analysis, we found that overexpression of miR-21, like ABC, suppressed lipid accumulation in response to a high cholesterol diet and increased accumulation of acylcarnitines.

**Conclusions:** Here we characterize microRNA dysregulation in MASH and MASH-HCC in patients, identify miR-21 as increasingly dysregulated from MASH to MASH-HCC, and delineate the impacts of miR-21 overexpression on lipid metabolism and hepatocarcinogenesis in zebrafish β-catenin-driven HCC. This study shows that miR-21, which is similarly dysregulated in human and zebrafish HCC, promotes lipid metabolic changes that may help drive hepatocarcinogenesis.

## Introduction

Hepatocellular carcinoma (HCC) is the third-leading cause of cancer-related death globally and maintains a median 5 year survival of just 16%^1,2^. HCC incidence is increasing in parallel with the growing prevalence of obesity and related risk factors including dyslipidemia, type 2 diabetes, and the proinflammatory state that accompanies the metabolic syndrome. These metabolic dysregulations are associated with the development of metabolic dysfunction-associated steatotic liver disease (MASLD)^3^. Advanced stages of MASLD (metabolic dysfunction-associated steatohepatitis, MASH) are characterized by inflammation, fibrosis, and the accumulation of lipotoxic lipid species such as acylcarnitines^4^, ceramides^5^, and unesterified cholesterol^6^, culminating in cirrhosis, which is associated with a 2.6% yearly cumulative HCC incidence (MASH-driven HCC, MASH-HCC)^7^. The molecular mechanisms driving the transition from MASH to MASH-HCC are incompletely understood^8^.

Mutations leading to the stabilization of the Wnt pathway co-transcriptional activator β-catenin (encoded by *CTNNB1*) are among the most frequent oncogenic events in MASH-HCC, occurring in approximately 30% of tumors^9,10^. Activated β-catenin (ABC) increases fatty acid oxidation and glutamate metabolism^11–13^.

Transgenic zebrafish expressing hepatocyte-specific ABC develop HCC with ∼80% penetrance as adults; zebrafish ABC-HCC is morphologically, transcriptomically, and metabolically similar to human HCC^14,15^. ABC zebrafish show robust liver enlargement by 6 days of age due to hepatocyte hyperproliferation, providing a facile platform for testing the effects of drugs and genetic manipulations on an HCC-related phenotype^14,16^.

We previously reported that ABC causes significant changes to acylcarnitines and triglycerides in cultured human liver cancer cells and in zebrafish liver^11,17^. ABC also promotes the oxidation of triglycerides and regulation of Ppara signaling^11^ in mice. Inhibition of fatty acid oxidation (FAO) pharmacologically in mice blunts HCC growth^11^. Lipidomics results from patient samples suggest that there is an increase in FAO during the transition from MASH to MASH-HCC^18^. Together these studies support the hypothesis that ABC promotes FAO to drive HCC, but the mechanism underlying ABC’s effects on FAO and other aspects of lipid metabolism are not well defined.

MicroRNAs (miRs) are 20-22 nucleotide RNA molecules that regulate the transcriptome^19^ by promoting mRNA degradation through direct interaction of conserved sequences between the miR and the 3’UTR of mRNA. miR targets can number in the 10s to the 100s^20^, and both direct and downstream effects of miRs can cause changes to gene expression^21,22^. Several miRs including miR-21^23,24^, miR-122^25^, miR-33^26^, and others^27^ have been shown to be dysregulated in serum and/or liver tissue of MASLD-HCC or HCC patients. The liver readily takes up oligonucleotides, facilitating hepatic delivery of miR mimics and antagonists such as miR-122 inhibitors, which safely and effectively reduce hepatitis C virus infection in the clinic^25,28^. Thus, miRs are enticing targets for HCC therapy.

Here we identify miR-21 expression as increasingly dysregulated from normal liver to MASH to MASH-HCC in patient tissues and in zebrafish ABC-HCC. We show that overexpression (OE) of hepatocyte-specific miR-21 enhanced ABC-driven larval liver enlargement and promoted HCC in adult zebrafish. We found that miR-21OE and ABC resulted in similar changes to hepatic steatosis in response to a high cholesterol diet. We reveal that miR-21OE drove HCC-associated lipid metabolic changes, providing insights into mechanisms of MASH-HCC.

## Results

### Identifying conserved dysregulated miRNAs in human and zebrafish HCC

To identify dysregulated, therapeutically relevant miRNAs in MASH and MASH-HCC, we utilized Nanostring to analyze patient samples from the University of Utah Pathology Archives (n = 4 or 7 for each group)(Supplemental Table 1). Out of the 827 miRNAs interrogated by Nanostring/nSolver, 28 miRNAs were significantly altered in MASH cirrhosis compared to non-cirrhotic control samples (14 up and 14 down) and 31 miRNAs were significantly altered in MASH-HCC compared to adjacent non-tumor (20 up and 11 down)(Supplemental Table 2). Four miRs—*let-7f, miR-15b, miR-21,* and *miR-32*—were significantly upregulated in both MASH and MASH-HCC, suggesting progressive upregulation with advancing liver disease (Table 1).

**Table 1.**
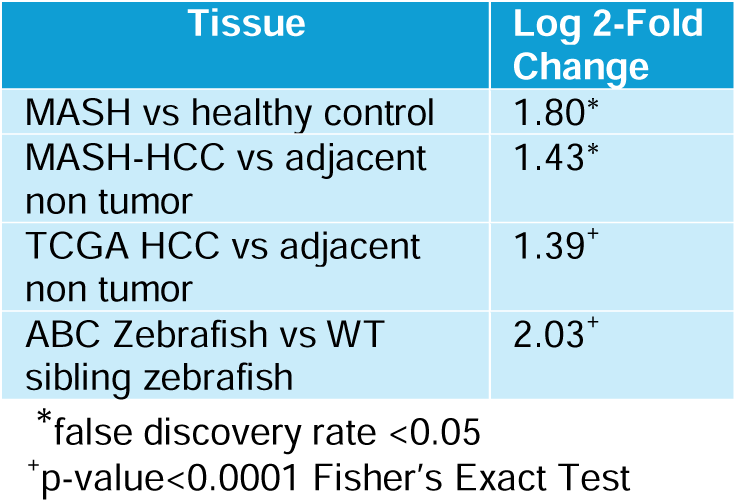
miR-21 levels in MASH, MASH-HCC, TCGA HCC, and transgenic zebrafish models of HCC.

We next validated our findings utilizing miRNA data from The Cancer Genome Atlas Liver and Hepatocellular Carcinoma (TCGA- LIHC) database^9^. We removed any patient samples that did not have paired sequencing, had a diagnosis other than or in addition to HCC, or had prior treatment, leaving 45 patients for analysis. Out of 623 miRNAs, 303 were significantly dysregulated (184 up and 119 down), including *miR-21* (Table 1, Supplemental Table 3).

To identify miRs with a conserved role in hepatocarcinogenesis across species and focus on those that might mediate the effects of ABC, we profiled miRs in zebrafish ABC-HCC. We performed miRNA sequencing and DESeq analysis of ABC-HCC and wildtype non-transgenic control siblings (WT)(n = 5 per group). Out of 212 detected pre-miRs, 82 were significantly dysregulated (43 up and 39 down regulated), including dre-miR-21-1 (L2FC 2.0, padj=1.5e-14) and dre-miR-21-2 (L2FC 1.8, padj=8.7e-7)(Supplemental Table 4). We used miRbase to identify 100 homologous miRNAs present in the 827 human miRs interrogated by Nanostring and in the 212 dre-pre-miRs identified by miR-seq. Of these homologous miRNAs, 16—including miR- 21—were significantly dysregulated in both humans and zebrafish, a significant overlap (Table 2, Figure 1A)(Supplemental Table 5). Together our analyses identify miR- 21 as one of the most robustly upregulated miRNAs in MASH- HCC and suggest shared mechanisms of miRNA-based modulation of HCC in zebrafish and humans (Table 2).

**Figure 1.**
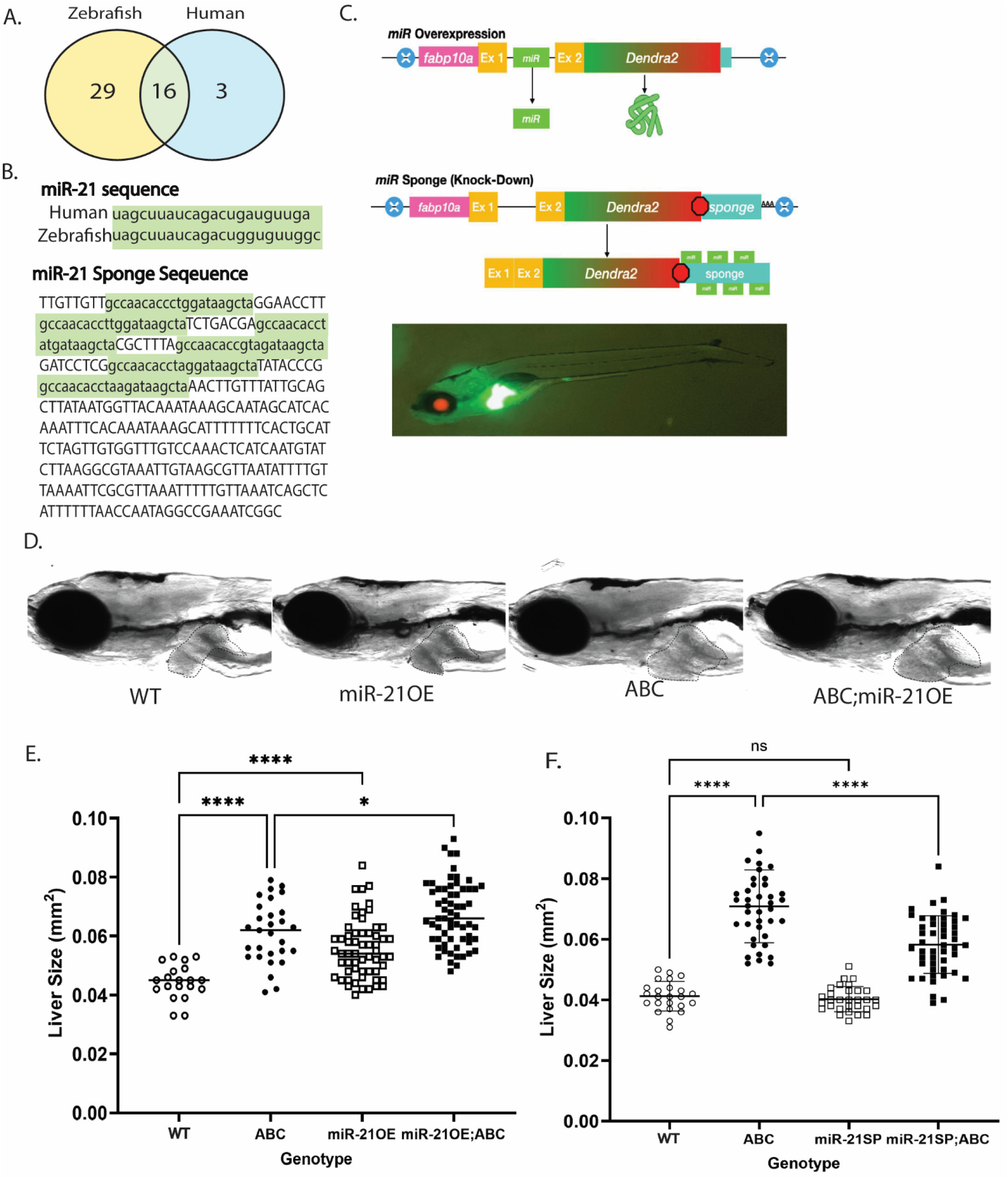
miR-21 overexpression (miR-21OE) or sponge (mir-21SP) alters zebrafish larval liver size compared to non-transgenic wildtype siblings (WT). A. Overlap of significantly altered miRs in zebrafish expressing activated β-catenin (ABC)(yellow) and human HCC samples (blue). B. Comparison of human miR- 21 sequence to zebrafish sequence and specific sequences used for overexpressing and sponging miR-21. C. Schematic of plasmid constructs and image of zebrafish larva with fluorescent red eyes and green liver indicating transgene expression. D. Representative brightfield images depicting larval zebrafish livers at 6 dpf. E and F. Liver size at 6 dpf comparing miR-21OE (E) and miR-21SP (F) to WT in the presence and absence of ABC. P values determined with GraphPad Prism, one-way ANOVA: ns, not significant; *, p < 0.05; ****, < 0.001.

**Table 2.** Homologous miRs dysregulated in HCC.

|  |  | miRNAs in human<sup>+</sup> |  |
| --- | --- | --- | --- |
|  |  | Sig | NS |
| miRNAs in zebrafish<sup>++</sup> | Sig | 16*** | 29 |
|  | NS | 3 | 52 |

<sup>+</sup>Sig, significant (p<0.05) as determined by Nanostring nCounter Human v3 miRNA Expression Assay kit and nSolver v4.0.70 software; NS, not significant.
<sup>+</sup>Sig, significant (p<0.05) as determined by DESeq; NS, not significant.
\*\*\*Fisher's Exact p<0.001, comparing overlap of significantly dysregulated homologous miRNAs in zebrafish and human.

### Modeling dysregulated miRs in zebrafish HCC

To characterize the role of miR-21 in zebrafish hepatocarcinogenesis, we generated transgenic zebrafish lines that either overexpress (miR-21OE*, Tg(fabp10a:Dendra2miR-21OE;cryaa:mCherry)*) or sponge (miR-21SP*, Tg(fabp10a:Dendra2miR-21SP;cryaa:mCherry)*) miR-21 under the control of the hepatocyte-specific *fabp10a* promoter^29^ (Figure 1B-C). We found that miR-21OE significantly increased larval liver size at 6 days post fertilization (6 dpf) from 0.045 mm^2^ to 0.056 mm^2^ (24% increase compared to wildtype non-transgenic control siblings (WT), p < 0.001)(Figure 1D-E). miR-21OE increased ABC-driven larval liver overgrowth from 0.061 mm^2^ to 0.067 mm^2^ (10% increase, p<0.05). Conversely, miR-21SP attenuated ABC-driven larval liver enlargement from 0.071 to 0.058 mm^2^ (18% decrease, p<0.0001)(Figure 1F). Histologic analysis of zebrafish livers at 12 months of age revealed that 11% of miR-21OE had HCC and 61.5% had mild changes (p<0.05 compared to WT)(Supplemental Figure 1). miR-21OE also led to HCC in the presence of loss-of-function mutation in the tumor suppressor *p53* (Supplemental Figure 1). These results support the hypothesis that miR-21 promotes HCC in zebrafish.

### miR-21OE, like ABC, suppresses hepatic lipid deposition in response to high cholesterol diet in larvae

To understand the role of miR-21 on hepatic lipid metabolism in zebrafish, including under conditions of metabolic stress, we fed miR-21OE larval zebrafish and wildtype non-transgenic control siblings (WT) a normal control diet (NCD) comprised of GEMMA Micro 75 zebrafish food or a high cholesterol diet (HCD) created by supplementing this commercial diet with 10% by weight cholesterol^30^ (Figure 2A). Steatosis was quantified by blinded examination of hematoxylin and eosin (H&E)-stained slides and by Oil red O (ORO) staining of whole-mounted larvae^31^. ORO stain highlights neutral lipid droplets that are predominantly comprised of triglycerides and cholesterol oleate^31^, and it has not been described to mark lipotoxic lipid species such as acylcarnitines^4^, ceramides^5^, and unesterified cholesterol^6^. In WT zebrafish, HCD increased steatosis assessed by H&E from 2% to 20% (p<0.0001) and increased ORO staining (p<0.001) (Figure 2B-D). miR-21OE suppressed HCD-induced steatosis, decreasing it from 20% to 6% by H&E (p<0.001) and decreasing ORO staining (p<0.0001)(Figure 2B-D). ABC-HCC zebrafish also showed decreased ORO staining on HCD compared to WT (p<0.0001) (Figure 2B,C). Overall, miR-21OE suppressed fat droplet accumulation in zebrafish liver in a manner analogous to ABC, suggesting common effects on lipid metabolism by both genes.

**Figure 2.**
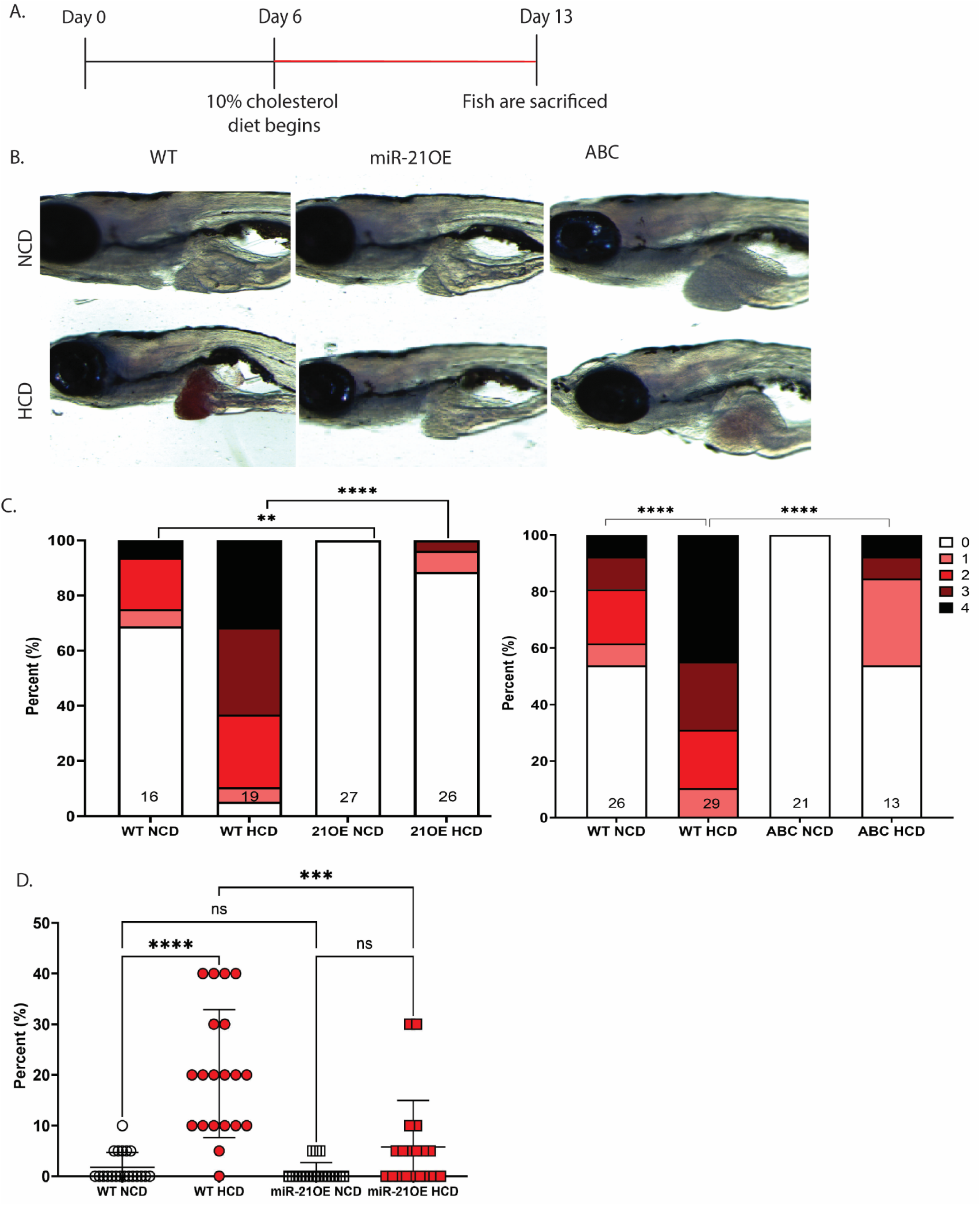
miR-21OE decreases steatosis in larval zebrafish fed a high cholesterol diet. A. Timeline of diet treatment. B. Representative brightfield images of 13 dpf miR-21OE (21OE), ABC, and non-transgenic wild-type control (WT) sibling zebrafish after Oil Red O staining (ORO). C. Quantification of ORO of larval liver images. Scoring is from 0-4. N values are indicated at the bottom of each column. P values determined with GraphPad Prism, Kruskal-Wallis test: ns; not significant; ****, p < 0.0001. D. Quantification of steatosis by hematoxylin and eosin staining. P values determined using GraphPad Prism, one-way ANOVA: ns, not significant; ***, p < 0.001, ****, p < 0.0001.

### miR-21OE downregulates expression of lipid metabolic genes

To identify potential targets of miR-21, including those dependent on diet, we performed RNA sequencing of livers dissected from 13-day-old miR-21OE zebrafish and non-transgenic wildtype control siblings (WT) on normal control diet (NCD) or high cholesterol diet (HCD) (Figure 3A-D, Supplementary Figure 3, and Supplementary Table 6a-d). Overall, 181 HCD-altered genes (107 down and 74 up) were significantly dysregulated when comparing WT-HCD to WT-NCD (Supplemental Table 6a), of which 12 were related to cholesterol metabolism (GO:0008203, FDR=1.71E-10) or biosynthetic process (GO:0006695, FDR=2.67E-08), confirming that HCD alters cholesterol metabolism (Figure 3E).

**Figure 3.**
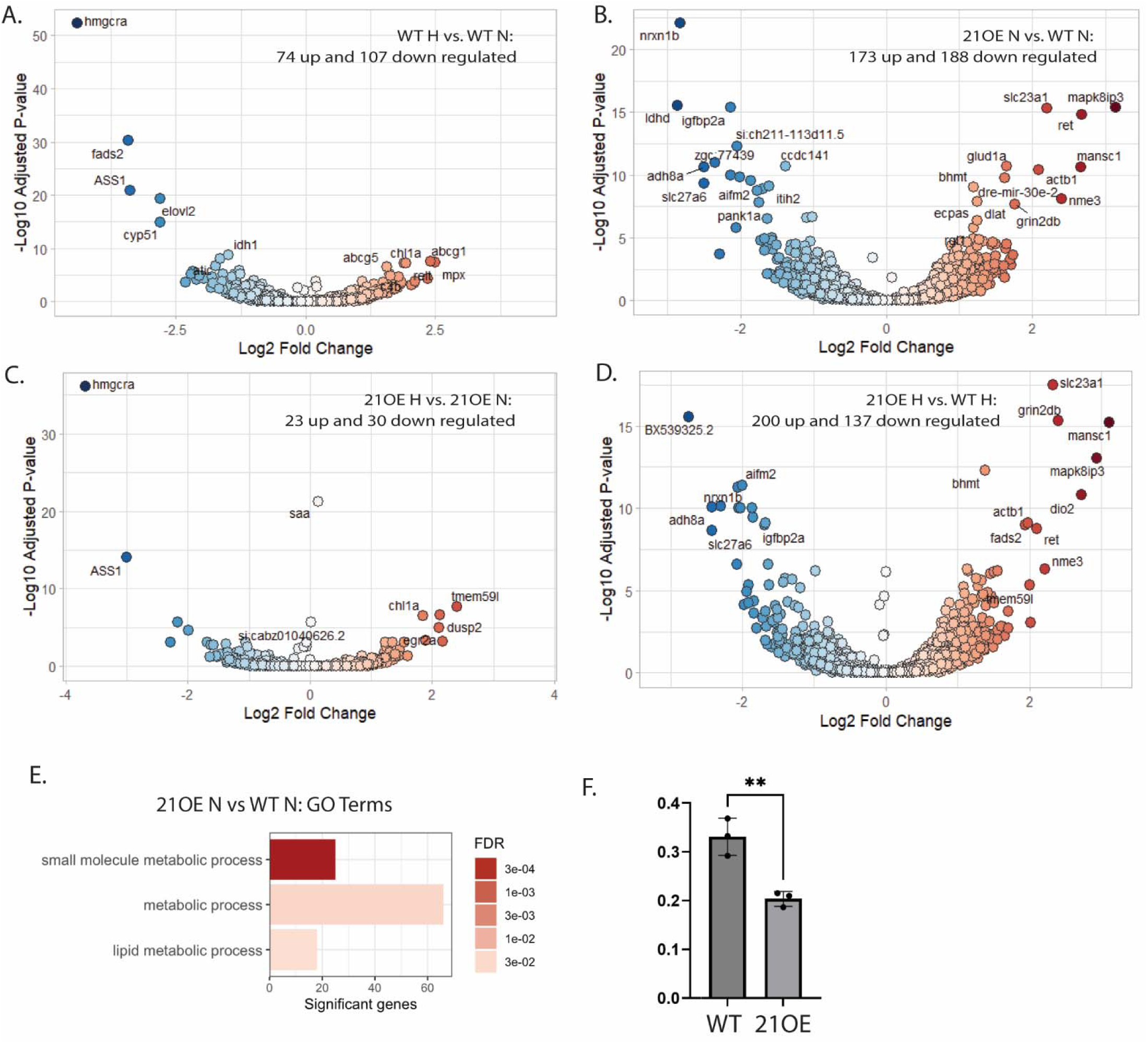
Gene expression changes induced by miR-21OE and/or high cholesterol diet in larval zebrafish. A-D. Volcano plots generated from RNA-seq data of miR-21OE (21OE) and/or non-transgenic wildtype control sibling (WT) zebrafish livers of larvae (13 dpf) administered normal control diet (N) or high cholesterol diet (H). E. Significant gene ontology (GO) terms for mir-21OE compared to WT on normal control diet, RNA-seq data. F. qPCR comparing NPC2 in WT and miR-21 OE livers for zebrafish larvae (13 dpf) on a normal control diet. P value determined using GraphPad Prism, Student’s t-test: **, p < 0.01.

We found that miR-21OE zebrafish had significantly higher levels of dre-mir-21-1 compared to controls (padj<0.01), confirming overexpression of miR-21 through the transgene construct (Supplemental Table 6b).

We identified 188 genes that were significantly downregulated in miR-21OE zebrafish compared to WT on NCD, 137 genes that were significantly downregulated in miR-21OE zebrafish compared to WT on HCD, and 70 genes that were significantly downregulated in both conditions (Supplemental Table 6b-c).

The miR-21OE-altered genes on NCD were enriched for small molecule metabolic process, metabolic process, and lipid metabolic process (Figure 3E). There was also a strong enrichment for genes involved in TCA cycle and glutamate metabolism like *mdh2, pdk1, got2a, slc1a3a, grin2db,* and *glud1a* (Supplemental Table 6b).

Downregulated genes in miR-21OE compared to WT on HCD included those involved in cholesterol metabolism and biosynthetic process (Supplemental Table 6c). Many downregulated genes contained one or more miR-21 seed sequences in their 3’UTRs according to our manual review of the UCSC Genome Browser and/or TargetScan^32,33,34,35^ (Table 3). Some of these genes, including *hmcgra*^36^, *ggcx*^37^, and *igfbp3*^38^, are established miR-21 targets in humans. We validated that at least six of these genes, including *npc2*, were significantly decreased with miR-21OE by performing qPCR on independent samples (Figure 3F and Supplemental Figure 2B).

**Table 3.**
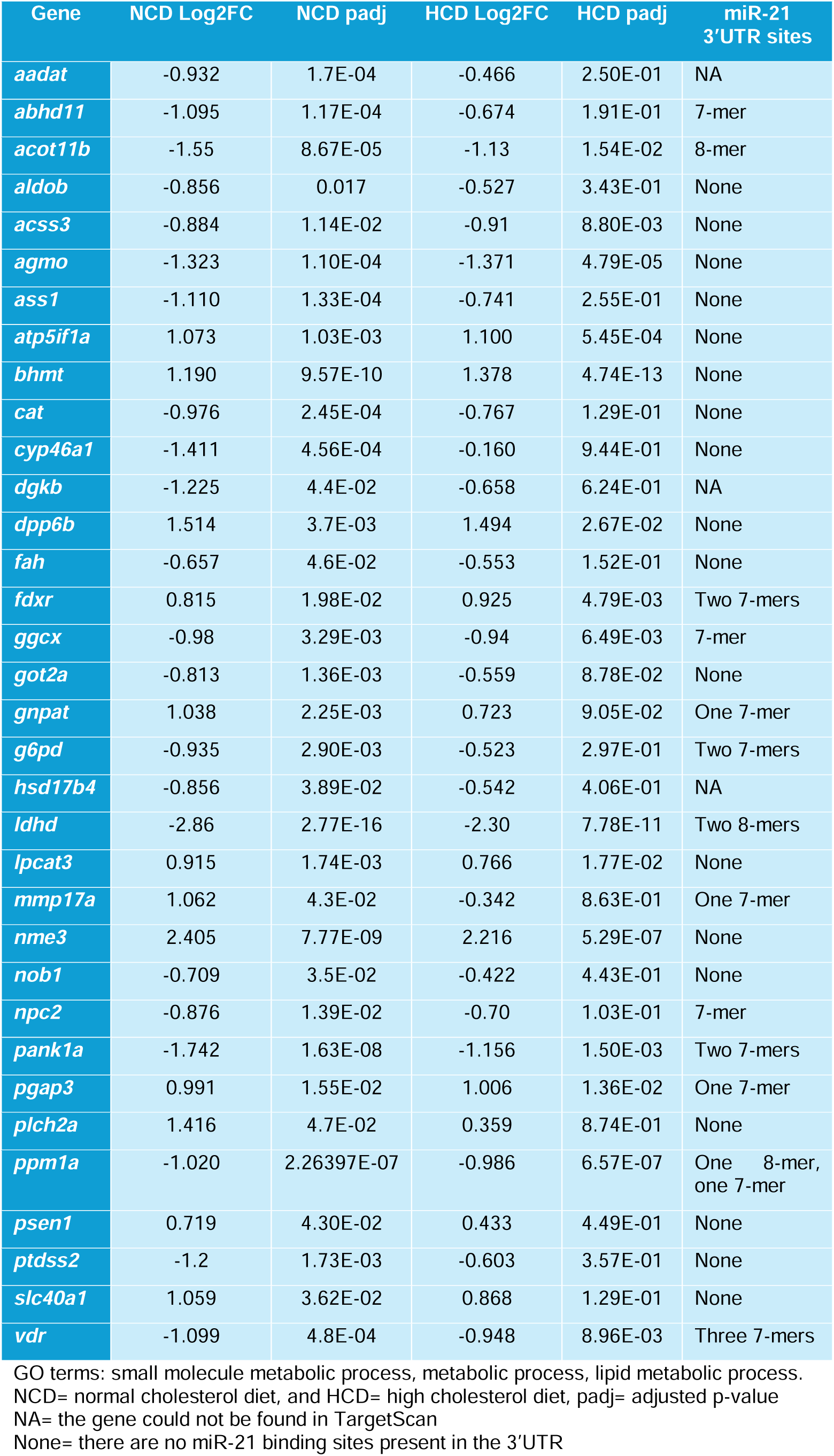
Metabolic genes altered by miR-21OE in 13 dpf larvae.

### miR-21OE, like ABC, suppresses hepatic lipid deposition in response to high cholesterol diet in adults

To further define the effects of miR-21OE and ABC on lipid metabolism, we administered 10% high cholesterol diet (HCD) or normal control diet (NCD) to adult zebrafish (Figure 4A). In WT zebrafish, HCD increased hepatic lipid deposition as quantified by ORO staining (p<0.0001) but was not sufficient to cause HCC (Figure 4B-D). Similar to larval zebrafish, adult ABC and miR-21OE zebrafish on HCD showed reduced lipid deposition compared to WT on HCD (p<0.0001 and p<0.0001, Figure 4B-D). On NCD, male and female ABC zebrafish had significantly higher HCC incidence (69% and 11%) compared to wildtype non-transgenic control siblings (WT)(0% and 0%, p<0.0001 and p<0.001)(Figure 4D), as we reported previously^14,15^. HCC incidence was not significantly altered by HCD (Figure 4D) in any genotype or sex. miR-21OE zebrafish rarely showed HCC while HCC was never observed in WT, though this difference was not statistically significant (Figure 4D). Zebrafish overexpressing both ABC and miR-21OE showed less HCC than ABC alone, though this difference was not statistically significant (Figure 4D). These data demonstrate that miR-21OE, like oncogenic ABC, suppresses normal lipid droplet accumulation in response to high cholesterol diet.

**Figure 4.**
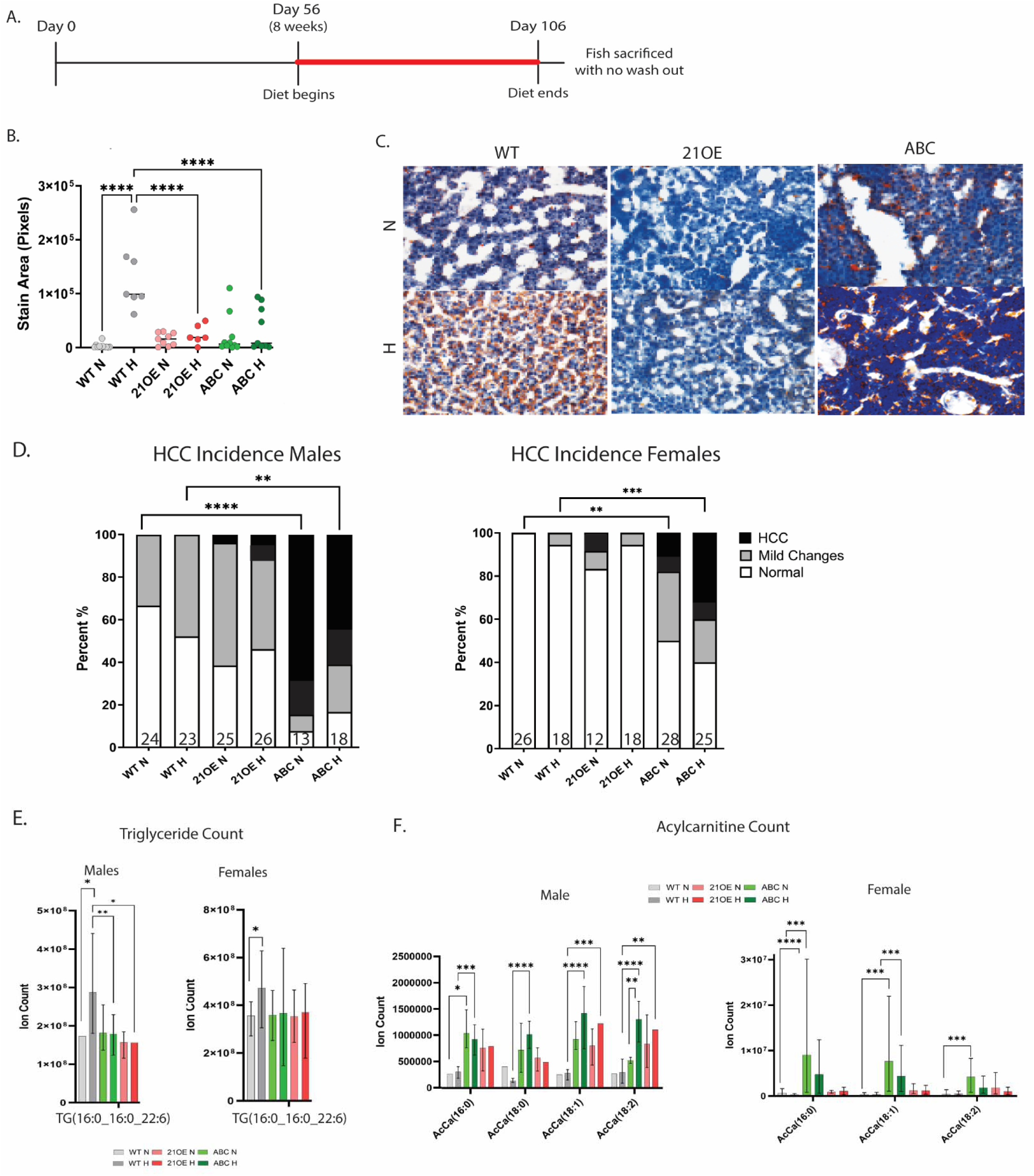
miR-21OE and ABC decrease steatosis, triglycerides, and acylcarnitines in adult zebrafish on a high cholesterol diet. A Timeline for feeding high cholesterol diet (H) or normal control diet (N) to adult zebrafish. B. Quantification of ORO for miR-21OE (21OE), ABC, and non-transgenic wild-type control (WT) sibling zebrafish livers. C. Representative ORO images. D. Histology assessed by hematoxylin and eosin (H&E) stain. N values indicated at bottom of column. P values determined with GraphPad Prism, one-way ANOVA. Comparing 21OE to WT and 21OE to ABC on either diet were not significant. E. Most abundant triglyceride counts. F. Most abundant acylcarnitine counts. Two-way ANOVA. For D-F: **, p < 0.01, ***, p < 0.001, ****, p < 0.0001.

### miR-21OE, like ABC, leads to decreased triglycerides and increased acyl carnitines

We next analyzed the changes to the lipidome in adult miR-21OE, ABC, and WT zebrafish administered a normal control (NCD) or high cholesterol diet (HCD)(Supplemental Figure 3). The most striking diet-induced changes in WT zebrafish fed HCD were in triglycerides (TG): the most abundant TG (16:0_16:0_22:6), increased in both males (p<0.05) and females (p<0.05)(Figure 4E and Supplemental Figure 4). Both ABC and miR-21OE suppressed the HCD-induced increase in triglycerides, though this effect was only statistically significant in males (Figure 4E and Supplemental Figure 3).

In ABC fish, the largest changes were to acylcarnitines (AcCa). Male (M) and female (F) ABC zebrafish showed a significant increase in AcCa (16:0) on both NCD and HCD (p<0.05 (M-NCD), p<0.001 (M-HCD), p<0.0001 (F-NCD), p<0.001 (F-HCD)) (Figure 4F). Other abundant AcCa species including AcCa (18:0), AcCa (18:1), and AcCa (18:2) were also significantly increased by ABC. miR-21OE also tended to increase AcCa compared to WT, though this effect was only statistically significant for AcCa (18:1)(p<0.0001) and AcCa (18:2)(p<0.001) in males on HCD (Figure 4F and Supplemental Figure 3).

## Discussion

Here we show that miR-21 is upregulated in liver tissue from MASH and MASH-HCC patients and in zebrafish ABC-HCC, supporting a conserved role for miR-21 in hepatocarcinogenesis. Our findings in MASH patients confirm the findings of Rodrigues et al., who found that miR-21 levels increased in MASLD and MASH in a European cohort^18^. Our finding that miR-21 is also upregulated in MASH-HCC compared to adjacent non-tumor tissue, together with prior results indicating a positive correlation between miR-21 levels and age^18^, suggests progressive dysregulation of miR-21 during MASH-driven hepatocarcinogenesis.

We found that miR-21 overexpression was sufficient to promote zebrafish larval liver overgrowth and enhance ABC-driven larval liver enlargement. miR-21 overexpression increased HCC in adult zebrafish, albeit to a lesser extent than ABC. Sponging miR-21 suppressed ABC-driven larval liver enlargement but did not affect larval liver size in the absence of ABC. We discovered that overexpression of miR-21 in zebrafish liver caused significant changes to genes involved in lipid and glutamate metabolism, which are also dysregulated by ABC^11,12,39^. In human umbilical vein endothelial cells, infection with ABC increases miR-21 expression^40^, while in glioma cells siRNA targeting ABC decreases miR-21 expression^40^. Together these findings suggest that miR-21 may be a direct or indirect downstream target of ABC that mediates some of the effects of ABC during tumorigenesis.

To understand the intersection of HCC, diet, ABC, and miR-21 we administered zebrafish a 10% high cholesterol diet (HCD), which induces hepatic steatosis, immune infiltration, and hepatocyte enlargement^30,41–43^. Excess cholesterol induces changes to lipid metabolism in the hepatocyte through altered regulation of SREBP2^44,45^, SREBP1c^46^, and the assortment of PPAR transcription factors^47^. We found that HCD led to downregulation of *ldlra, hmgcra*, and *fads2,* (Supplemental table 5A and B) which are involved in de novo cholesterol synthesis^48^, cholesterol homeostasis^49^, and the regulation of polyunsaturated fatty acids^50^, respectively. There was also downregulation of *srebf2* and *pparaa* in the WT and/or miR-21OE livers on HCD. HCD caused increases to *stard4* and *apoa4*, which are involved in cholesterol esterification^51^ and triglyceride export^52^ and mediate cross talk between cholesterol and lipid metabolism through srebp1c^53^. miR-21OE zebrafish fed HCD showed changes to these genes in the same direction as non-transgenic wildtype control zebrafish but had a less dramatic decrease in *fads2* and a greater increase in *apoa4*. This finding suggests a mechanism in which miR-21OE zebrafish may suppress hepatic lipid accumulation, as *fads2* and *apoa4* are involved in triglyceride accumulation^52,54^.

When comparing genes that were significantly altered in miR-21 on HCD versus NCD, we found enrichment of GO terms small molecule metabolic process (GO: 0044281) and organophosphate synthetic process (GO: 0090407). These processes were not significantly altered in WT on the HCD diet, suggesting that miR-21OE may rewire metabolism such that hepatocytes avoid excessive lipid droplet formation in response to excess cholesterol. This hypothesis is corroborated by our ORO and H&E staining results showing that miR-21OE decreased accumulation of lipid droplets and triglycerides in response to a high cholesterol diet. Multiple lines of evidence suggest that suppression of the accumulation of triglycerides in lipid droplets represents a pathologic, tumor-promoting response. First, we noted similar suppression of lipid accumulation with overexpression of ABC, a well-established hepatic oncogene^13,56^. Second, in human^56^ and mouse^57^ MASH, steatosis decreases at late stages when HCC rates are highest. Decreased steatosis during progression from MASH to HCC may reflect alterations in fatty acid metabolism, an emerging hallmark of cancer^58^.

We found that acyl carnitines were the most substantially altered lipid species in response to ABC and miR-21 overexpression, confirming our prior results in zebrafish and human cell lines^17^. Prior studies in mouse osteoblasts^39^ and mouse HCC^11^ have also shown that ABC promotes fatty acid oxidation. Serum levels of long-chain acyl carnitines increase during progression from MASLD to MASH-HCC^59,60^ and are elevated in ABC zebrafish compared to controls^17^. Our current findings support a positive correlation between acylcarnitine levels and tumor burden. ABC fish had the highest levels of acylcarnitines and the most HCC, miR-21OE had intermediate levels of acylcarnitines and occasional HCC, and WT had the lowest acylcarnitine levels and no HCC. Together these data suggest that like ABC, miR-21 promotes fatty acid oxidation, potentially resulting in a lipotoxic state that can drive further HCC progression^61^ and support tumor burden.

Across many cancer types, miR-21 is one of the most well-established oncogenic miRNAs (“onco- miRs”)^62–64^. Nonetheless, prior HCC studies have yielded seemingly conflicting results regarding whether miR- 21 is pro- or anti-tumorigenic in the liver or if the role of miR-21 changes as disease progresses. On one hand, hepatic miR-21 levels are elevated in MASH-HCC^18^, and miR-21 knockout decreases liver tumorigenesis in mice fed a choline-deficient diet^18^. On the other hand, in some situations it appears that miR-21 may be protective against HCC: When HFD is combined with the carcinogen diethylnitrosamine (DEN) given at 3 weeks, constitutive miR-21 knockdown leads to increased liver tumor burden while miR-21 mimic decreases liver tumors^65^. Constitutive or hepatocyte-specific miR-21 knockout leads to increased tumorigenesis following DEN or in liver-specific PTEN knockout mice^66^.

This study examining the effects of miR-21OE at various time points and under different dietary conditions provides some insights into this apparent discrepancy. We propose that in the setting of a non-neoplastic liver, such as with mice fed a choline-deficient diet, adult zebrafish that are not overexpressing another oncogene (Figure 4, 21OE vs WT groups), or larval zebrafish with β-catenin-driven hepatocyte hyperproliferation (Figure 1), miR-21 overexpression drives lipid dysregulation and HCC. However, once HCC has been established or induced via genetic alterations and/or carcinogen treatment^65,66^, such as in the setting of ABC adult zebrafish (Figure 4D, ABC;21OE vs ABC groups), miR-21-driven metabolic dysregulation decreases tumor burden by promoting tumor cell death. This hypothesis is supported by prior reports that miR- 21 knockout decreases TUNEL staining and active caspase-2 levels in mice fed a methionine-choline deficient diet^67^. Future work in our laboratory will focus on defining the link between miR-21, metabolism, cell death, and liver tumorigenesis.

## Methods

### miRNA expression analysis in University of Utah patient samples

We searched the University of Utah Pathology Archives and identified samples from 7 patients with MASH-HCC, 4 patients with MASH (and no clinical or pathologic evidence of HCC), and 4 control patients without cirrhosis or MASH (Supplemental Table 1). Patients were excluded if they had a history of infection with hepatis B or C virus or significant alcohol use. The University of Utah High-Throughput Genomics Shared Resource (HTG) Core extracted RNA from paraffin-embedded, formalin-fixed liver tissues using the Qiagen miRNeasy FFPE kit and assessed miRNAs using the Nanostring nCounter Human v3 miRNA Expression Assay kit (CSO-MIR3-12). miRNA counts were analyzed using nSolver v4.0.70. miRNAs were considered significantly dysregulated with a log2foldchange +/- 0.5 and FDR <0.05.

### miRNA expression analysis in TCGA patient samples

To determine significantly dysregulated miRNAs in patients with HCC, human paired tumor and non-tumor miRNA expression quantification normalized count data was gathered from The Cancer Genome Atlas Liver Hepatocellular Carcinoma (TCGA-LIHC) GDC 23.0 Data Release using TCGA biolinks package^68^.

Patient samples were excluded from analysis if history indicated they had received treatment prior to biopsy or had a final diagnosis of a malignancy other than, or in addition to, HCC. In total, 45 patients were eligible for analysis. Differentially expressed miRNAs were identified with DESeq2 version 1.34.0^69^. miRNAs were considered significantly dysregulated with a log2foldchange +/- 0.5 and padj <0.05.

### Zebrafish husbandry

Zebrafish (Danio rerio) lines were maintained under standard conditions^70^. In brief, embryos and larvae were cultured in egg water (2.33 g Instant Ocean in 1 L Milli-Q water with 0.5 ml Methylene Blue) or low-salt egg water (60 mg Instant Ocean in 1 L Milli-Q water) and incubated at 28.5°C. At ∼5 days post fertilization (dpf), zebrafish were transitioned to a recirculating system. Juvenile and adult zebrafish were fed brine shrimp, commercial powdered food, GEMMA, and/or commercial powdered food, and housed on a recirculating system. In addition to wildtype AB zebrafish (WT), we also used previously established *Tg(fabp10a:pt-*β*-cat)* zebrafish, which express hepatocyte-specific activated β-catenin (ABC)^14^.

### miRNA expression analysis in zebrafish livers

Five transgenic male ABC zebrafish and five non-transgenic male wildtype control siblings (WT) were raised under standard conditions to adulthood as described above and sacrificed at 4 months of age. Livers were dissected from the body cavity, a small portion of each liver was fixed in 4% PFA and submitted to ARUP Research Histology to generate H&E-stained slides, and the remaining liver was snap frozen. H&E-stained slides were examined by a pathologist (K.J.E.) to confirm the diagnosis of HCC (ABC) or no significant pathologic abnormalities (WT). The HTG Core extracted RNA, prepared libraries using NEBNext Multiplex Small RNA library Prep Set, and sequenced using Illumina HiSeq 50 bp single read sequencing. miRNAs were aligned to GRCz10 and counted using miRbase. Differentially-expressed miRNAs were identified with DESeq2 version 1.34.0^69^. miRNAs were considered significantly dysregulated with a L2FC +/- 0.5 and padj <0.05.

### Generation of transgenic zebrafish to overexpress or sponge miR-21 (miR-21OE and miR-21SP)

To generate the *fabp10a:miR-21* plasmid (*fabp10a-miR-21-1-Dendra2,cryaa:mCherry*), we first made *miR-21-1-Dendra2* by amplifying *dre-mir-21-1* from AB zebrafish genomic DNA using primers with BbsI cut sites (Supplemental Table 7) and inserting it into *Tol2-lyzC-Vector-Dendra2* (Addgene plasmid # 97101, a kind gift from Qing Deng)^71^. From this *Tol2-lyzC-miR-21-1-Dendra2* plasmid, we amplified *miR-21-1-Dendra2* with flanking XhoI cut sites and placed it downstream of the *fabp10a* promoter^72^ into an *I-SceI* meganuclease vector^73^ that also contained *cryaa:mCherry*^74,75^.

To generate the *fabp10a:miR-21SP* plasmid (*fabp10a-miR-21SP-Dendra2*), as outlined by Zhou et al. 2018^76^, we ordered gBlocks^TM^ (IDT, Coralville, Iowa) which contained 6 bulging miRNA binding sites with MfeI and BamHI cut sites at the 5’ and 3’ ends respectively (Supplemental Table 8). gBlocks and *Tol2-lyzC-Vector- Dendra2* were separately digested with MfeI and BamHI, gel extracted, column purified, and ligated to generate *Tol2-lyzC-miR-21SP-Dendra2.* From this *Tol2-lyzC-miR-21SP-Dendra2* plasmid, we amplified *miR- 21SP* with flanking XhoI cut sites and placed it downstream of the *fabp10a* promoter^72^ into an *I-SceI* meganuclease vector^73^ that also contained *cryaa:mCherry*^74,75^.

One-cell-stage embryos were microinjected with *fabp10a:miR-21* or *fabp10a:miR-21SP* plasmid, *I-SceI* meganuclease, *I-SceI* buffer, and Phenol Red as previously described^72^. Injected embryos with red eyes and green livers at 2–5 days post fertilization (dpf) were raised to adulthood and crossed to detect founders with germline transmission. We identified three unique miR-21OE founders, which all had similar phenotypes, and two unique miR-21SP founders, which all had similar phenotypes. The miR-21OE and miR-21SP lines were maintained by outcrossing them to wildtype AB zebrafish each generation. Transgenic zebrafish were distinguished from non-transgenic control siblings by the presence of red eyes and green livers at 3 dpf or later. Phenotypes were consistently seen starting at the F1 generation, and all published experiments were performed on F2 generation or later.

### Zebrafish diet studies

For zebrafish larval diet studies, zebrafish were maintained under standard conditions (egg water or low-salt egg water with no feeding) until 6 dpf. From 6 to 12 dpf, zebrafish were fed daily with GEMMA Micro 75 (normal control diet, NCD) or with GEMMA Micro 75 containing 10% by weight cholesterol (C8667, Millipore Sigma), prepared as previously described (high cholesterol diet, HCD)^30^. Zebrafish were maintained in 2.8L tanks. Water changes were performed daily to remove debris and exchange 1 L of low-salt egg water. At 13 dpf larvae were euthanized and fixed with 4% paraformaldehyde.

For zebrafish adult diet studies, zebrafish were maintained under standard conditions until 8 weeks of age. Beginning at 8 weeks of age, zebrafish were fed twice daily with GEMMA 75, 150, or 500 (normal control diet, NCD) or with GEMMA containing 10% by weight cholesterol, prepared as previously described (high cholesterol diet, HCD)^30^. Each zebrafish tank was also given 2 mL of suspended brine shrimp once a day.

Zebrafish were given the specialized diet for 50 days (7 weeks) and then sacrificed ∼12 hours after their last feeding.

### Quantification of liver size and steatosis

Quantification of larval liver size at 6 dpf was performed as previously described^77^. In brief, larvae were raised to 6 dpf under standard conditions, fixed in 4% paraformaldehyde (PFA), and dissected to remove the pectoral fins, cartilage, and skin to expose the liver and peritoneal cavity. Images were taken of each larva using a Leica dissecting microscope, and blinded image files were analyzed with FIJI/ImageJ. Fixed 13 dpf larvae were stained with Oil Red O (ORO) stain by washing 4% PFA-fixed fish in isopropanol for 30 minutes followed by the addition of fresh isopropanol with 0.3% dissolved ORO Stain. The fish were left to rock on orbital rocker for 90 minutes in this solution and washed once with isopropanol then PBS for 5 minutes each on orbital rocker^78^. Within 48 hours the larvae were dissected to expose the liver and imaged with a Leica dissecting microscope. Blinded images were given a semi-quantitative score based on relative ORO staining intensity on a scale from 0 (no staining) to 4 (complete, intense staining).

For adult zebrafish, frozen sections and ORO staining were performed by ARUP Research Histology.

Slides were blinded, and one representative image was taken of each slide using an Olympus BX53 microscope with Olympus DP73 camera and cellSens software. The area of red stain within each image was quantified with FIJI/ImageJ. One way ANOVA was used for statistical analysis.

### RNA sequencing and qPCR

Zebrafish at 13 dpf were euthanized, and livers were dissected and pooled (13 livers per sample) for RNA Extraction using PicoPure RNA extraction kit (Thermo, KIT0204). For sequencing, libraries were prepared by the HTG Core using NEBNext Ultra II Directional RNA Library Prep with rRNA Depletion Kit (Zebrafish).

Samples were barcoded, pooled, and sequenced using paired 150 bp sequencing on Illumina NovaSeq X. Reads were aligned to GRCv11 zebrafish genome. Genes with less than 10 counts were removed.

Differentially expressed transcripts were identified with DESeq2 version 1.40.2 and were considered significantly dysregulated with a log2foldchange +/- 0.5 and padj <0.05. Pathway analysis was completed using FishEnrichr and Gene Ontology Biological Process 2025. GO Terms graphs were made using ggplot R package.

Samples for qPCR were reverse transcribed using Super Script III kit (Catalog number 18080051).

Primer sequences are listed in Supplemental Table 9. qPCR master mixes were prepared consisting of 2.5% 100 μM forward primer, 2.5% 100 μM reverse primer, and 62.5% PowerTrack SYBR Green Master Mix (Thermo, A46109) in RNase-free water. Master mixes were combined 4:1 with the cDNA reactions and plated in duplicate. qPCR was performed using the LC480 PCR Lightcycler (Roche, 05015278001) using the “Mono Color Hydrolysis Probe/UPL probe” detection format. The temperature cycle consisted of an initial 2 min period at 95°C and 40 cycles of 95°C for 15 sec and 60°C for 50 sec set to single acquisition mode. The housekeeping gene β-actin was used as an internal control for cDNA quantification and normalization of amplified products. Data are reported as relative expression.

### Histological evaluation of zebrafish by H&E

Whole-body paraffin embedding, sectioning, and staining with hematoxylin and eosin (H&E) was performed by ARUP Research Histology. Sections were reviewed and scored by a board-certified pathologist (K.J.E.) in a blinded fashion using an Olympus BX53 microscope. For larval zebrafish (13 dpf), steatosis was scored by estimating the percentage of liver parenchyma involved by fat. For adult zebrafish, each zebrafish was assigned to one of the following categories as previously described^14^: (1) No changes, defined as no substantial cytological or architectural abnormalities; (2) Mild changes, defined as the presence of cytological abnormalities in the absence of substantial architectural abnormalities or vice versa; or (3) Hepatocellular carcinoma (HCC), defined as the presence of both architectural and cytological abnormalities. These terms were given a numerical score, 0-2, respectively, and then graphed and analyzed by one way ANOVA.

### Lipid Extraction

Adult zebrafish were grouped based on diet, sex, and genotype. Livers were isolated using a dissecting microscope and weighed. Livers were snap frozen in liquid nitrogen and placed on dry ice. For male samples, liver tissues from up to three animals of the same genotype/diet group were pooled to achieve a minimum of 15 mg. We analyzed 1-6 replicates per group.

10-30 mg of flash-frozen zebrafish liver tissue were isolated in 2 mL Safelock microcentrifuge tubes (Eppendorf, 022363352) with a 5/16 in. diameter stainless steel ball (Grainger, 4RJL8) chilled to -80°C. Tissue was homogenized at 25 Hz for 30 seconds under liquid nitrogen using a Retsch Cryomill (Retsch, 20.749.0001). Lipids were extracted by adding 1 mL of 75% methyl tert-butyl ether, 24% ul methanol, and 1% Splash Lipidomix Mass Spec Standard (Avanti Polar Lipids 330707) and incubating on ice for 15 minutes with intermittent vortexing, and phase separation was induced with 190 µl ultrapure water. The organic supernatant was transferred to a glass vial and dried with gaseous nitrogen. Analytes were resuspended in 25 µl/mg sample in 2:1:1 isopropyl alcohol:acetonitrile:water and transferred to amber mass spectrometry vials (Agilent, 5182-0716) with glass inserts.

### LC-MS Lipidomics

Lipid extracts were analyzed by LC-MS using a Vanquish HPLC system (Thermo Fisher Scientific) and a QExactive HF Orbitrap mass spectrometer (Thermo Fisher Scientific). Separation was achieved by C18 chromatography performed on an Acquity UPLC CSH C18 column (2.1 mm X 100 mm, 1.7 µm particular size, 130 Å pore size, Waters Co., 186005297). The chromatography gradient was formed by solvent A (10 mM ammonium formate and 0.1% formic acid in 60:40 acetonitrile:water) and solvent B (10 mM ammonium formate and 0.1% formic acid in 90:9:1 isopropyl alcohol:acetonitrile:water) at a constant flow rate of 350 µL/min. The gradient function was: 0 min, 30% B; 5 min, 43% B; 5.1 min, 50% B; 14 min, 70% B; 21 min, 99% B; 24 min, 99% B; 24.1 min, 30% B; 31 min, 30% B. Autosampler temperature was 4°C, column temperature was 30°C, and injection volume was 2 μL. Samples were injected into the mass spectrometer by electrospray ionization operating in positive ion mode. Lipid mass spectra were collected in full scan mode at 70,000 resolving power, and peaks were identified based on exact mass and retention times using El-MAVEN with comparison with known standards^79^.

## Statistical Methods of Analysis

GraphPad Prism V10.2 was used to perform statistical analyses and generate graphs. For larval liver size, liver to body ratio, percent steatosis, Brown-Forsythe and Welch ANOVA tests were performed. For differences in oil red o (ORO) staining and HCC diagnosis Mann-Whitney tests were performed. For lipidomics comparison a 2-Way ANOVA with Dunnett post hoc analysis was used for comparing genotype effects. To test diet effect across genotypes a 2-Way ANOVA with Sidaks post hoc analysis was used.

## List of Abbreviations

ABC: activated β catenin
HCC: hepatocellular carcinoma
MiR: microRNA
NCD: normal control diet
HCD: high cholesterol diet
AcCa: acylcarnitine
PE: phosphatidylethanolamines
PC: phosphotidylcholine
SM: sphingomyelin
TG: triglyceride
FAO: fatty acid oxidation

