## Supplemental Tables for "microRNA-21 promotes dysregulated lipid metabolism and hepatocellular carcinoma"

**Supplementary Table 1:** Clinical and demographic information for patients with MASH or MASH-HCC

| **Patient** | **Diagnosis Category** | **Age** | **Sex** | **Race** | **Ethnicity** | **Diagnosis (tumor)** | **Diagnosis (non-tumor)** | **Other clinical diagnoses** |
| --- | --- | --- | --- | --- | --- | --- | --- | --- |
| 1 | Cirrhosis | 37 | M | White | Non-Hispanic | No malignancy | Cirrhosis | Morbid obesity, MASH |
| 2 | Cirrhosis | 54 | M | Other | Hispanic/  Latino/a/x | No malignancy | Cirrhosis | DM, MASH |
| 3 | Cirrhosis | 44 | M | Other | Hispanic/  Latino/a/x | No malignancy | Cirrhosis with mild steatosis | MASH |
| 4 | Cirrhosis | 56 | F | Mexican American | Hispanic/  Latino/a/x | No malignancy | Cirrhosis | MASH |
| 5 | Non-cirrhotic | 68 | M | Other | Hispanic/  Latino/a/x | Metastatic adenocarcinoma | Non-cirrhotic | Metastatic colorectal adenocarcinoma |
| 6 | Non-cirrhotic | 69 | M | White | Non-Hispanic | Metastatic adenocarcinoma | Non-cirrhotic, mild steatosis | Metastatic colorectal adenocarcinoma |
| 7 | Non-cirrhotic | 69 | M | White | Non-Hispanic | Hemorrhagic cavernous hemangioma | Non-cirrhotic | Complex hepatic cysts, high cholesterol, DM |
| 8 | Non-cirrhotic | 61 | F | White | Non-Hispanic | Metastatic adenocarcinoma | Non-cirrhotic | Metastatic colorectal adenocarcinoma |
| 9 | HCC+matched non-tumor | 62 | M | White | Non-Hispanic | HCC | Cirrhosis | MASH |
| 10 | HCC+matched non-tumor | 74 | M | White | Non-Hispanic | HCC | Cirrhosis | MASH, DM |
| 11 | HCC+matched non-tumor | 68 | M | White | Non-Hispanic | HCC | Cirrhosis | MASH |
| 12 | HCC+matched non-tumor | 58 | M | White | Hispanic/  Latino/a/x | HCC | Cirrhosis | Cryptogenic cirrhosis, no significant alcohol history, obesity, high cholesterol |
| 13 | HCC+matched non-tumor | 69 | M | White | Unknown | HCC | Cirrhosis | MASH, obesity, OSA |
| 14 | HCC+matched non-tumor | 66 | M | White | Non-Hispanic | HCC | Cirrhosis | Probable MASH, no significant alcohol history, obesity |
| 15 | HCC+matched non-tumor | 68 | F | White | Non-Hispanic | HCC | Cirrhosis | MASH, obesity |

DM, Diabetes mellitus; OSA, obstructive sleep apnea

**Supplementary Table 2.** miRNAs are significantly dysregulated in MASH and MASH-HCC as identified by NanoString nCounter Human v3 miRNA Expression Assay kit.

| **Probe Name** | **MASH vs.**  **NC^+^** | | **HCC vs.** | |
| --- | --- | --- | --- | --- |
|  |  |  | **NT^++^** | |
|  | **L2Ratio** | **FDR** | **L2Ratio** | **FDR** |
| *hsa-let-7b-5p** |  |  | -0.92 | <0.001 |
| *hsa-let-7e-5p** | 2.07 | 0.04 |  |  |
| *hsa-let-7f-5p* | 2.06 | 0.01 | 2.39 | <0.001 |
| *hsa-miR-100-5p** |  |  | -1.45 | <0.001 |
| *hsa-miR-107** |  |  | 1.87 | 0.01 |
| *hsa-miR-10a-5p** | 2.17 | 0.05 |  |  |
| *hsa-miR-1246* | -2.77 | 0.01 |  |  |
| *hsa-miR-1285-5p* | -3.77 | <0.001 |  |  |
| *hsa-miR-132-3p* |  |  | 0.76 | 0.05 |
| *hsa-miR-145-5p* | 4.09 | 0.01 |  |  |
| *hsa-miR-146b-5p** | 2.24 | 0.02 |  |  |
| *hsa-miR-148b-3p* |  |  | 1.35 | <0.001 |
| *hsa-miR-151a-5p* |  |  | 0.9 | <0.001 |
| *hsa-miR-15a-5p** |  |  | 2.41 | 0.03 |
| *hsa-miR-15b-5p** | 1.84 | 0.05 | 3.01 | <0.001 |
| *hsa-miR-181a-5p* |  |  | -2.49 | <0.001 |
| *hsa-miR-186-5p* |  |  | -2.71 | 0.02 |
| *hsa-miR-193b-3p* |  |  | 1.21 | 0.01 |
| *hsa-miR-194-5p** |  |  | 3.17 | 0.04 |
| *hsa-miR-195-5p* | 2.43 | 0.01 |  |  |
| *hsa-miR-1972* | -3.54 | 0.02 |  |  |
| *hsa-miR-199a/b-3p* | 1.96 | <0.001 |  |  |
| *hsa-miR-200b-3p** | 2.39 | 0.04 |  |  |
| *hsa-miR-21-5p** | 1.8 | <0.001 | 1.43 | 0.01 |
| *hsa-miR-218-5p* | 1.15 | <0.001 |  |  |
| *hsa-miR-221-3p* |  |  | 0.64 | 0.03 |
| *hsa-miR-222-3p** |  |  | -1.18 | <0.001 |
| *hsa-miR-23a-3p** | 1.08 | 0.01 |  |  |
| *hsa-miR-24-3p** |  |  | 1.77 | <0.001 |
| *hsa-miR-28-5p* |  |  | 1.3 | 0.05 |
| *hsa-miR-30a-5p* |  |  | -3.22 | <0.001 |
| *hsa-miR-30b-5p* |  |  | 2.1 | 0.03 |
| *hsa-miR-30e-5p* |  |  | -2.84 | <0.001 |
| *hsa-miR-32-5p* | 1.7 | 0.02 | 2.49 | <0.001 |
| *hsa-miR-362-3p* |  |  | 0.52 | 0.04 |
| *hsa-miR-376a-3p* |  |  | -1.24 | <0.001 |
| *hsa-miR-379-5p* | -2.99 | 0.01 |  |  |
| *hsa-miR-421* | -4.81 | 0.03 |  |  |
| *hsa-miR-4443* | -3.62 | 0.02 |  |  |
| *hsa-miR-4488* | -2.83 | <0.001 |  |  |
| *hsa-miR-4516* | -3.41 | 0.02 |  |  |
| *hsa-miR-454-3p* |  |  | 0.58 | 0.04 |
| *hsa-miR-494-3p* |  |  | -7.35 | <0.001 |
| *hsa-miR-518b* |  |  | 1.01 | 0.03 |
| *hsa-miR-519d-3p* |  |  | 2.15 | 0.01 |
| *hsa-miR-548q* | -3.15 | 0.05 |  |  |
| *hsa-miR-575* | -5.7 | 0.02 | -4.38 | 0.05 |
| *hsa-miR-579-3p* |  |  | -4.1 | 0.04 |
| *hsa-miR-585-3p* | -2.95 | 0.04 |  |  |
| *hsa-miR-612* | -2.93 | 0.03 |  |  |
| *hsa-miR-630* | -7.49 | 0.01 |  |  |
| *hsa-miR-888-5p* | -3.16 | 0.02 |  |  |
| *hsa-miR-98-5p* |  |  | 1.6 | <0.001 |
| *hsa-miR-99b-5p** | 0.87 | 0.03 |  |  |

**^+^**MASH v NTNC: Comparing MASH samples to non-tumor non-cirrhotic samples from different patients

**^++^**HCC vs NT: Comparing MASH-HCC tumor sample to adjacent non-tumor in the same patients

* miRNA also significantly dysregulated in ABC zebrafish (Supplemental Table 5.2)

FDR: False discovery rate

**Supplementary Table 3.** 303 miRNAs significantly dysregulated in TCGA-LIHC

| **Gene** | **All Samples** | |
| --- | --- | --- |
|  | L2FC | padj |
| *hsa-mir-424* | -1.96 | 2.86E-52 |
| *hsa-mir-490* | -5.09 | 1.63E-41 |
| *hsa-mir-1258* | -4.00 | 1.80E-38 |
| *hsa-mir-130a* | -1.75 | 8.60E-38 |
| *hsa-mir-450b* | -1.57 | 3.62E-33 |
| *hsa-mir-139* | -1.93 | 2.53E-32 |
| *hsa-mir-675* | -2.68 | 2.03E-29 |
| *hsa-mir-450a-1* | -1.56 | 5.55E-23 |
| *hsa-mir-483* | -1.49 | 1.28E-22 |
| *hsa-mir-451a* | -1.75 | 1.35E-21 |
| *hsa-mir-199a-1* | -2.10 | 1.64E-20 |
| *hsa-mir-383* | -2.57 | 1.91E-20 |
| *hsa-mir-199a-2* | -2.09 | 2.70E-20 |
| *hsa-mir-542* | -1.16 | 1.68E-19 |
| *hsa-mir-144* | -1.54 | 3.82E-19 |
| *hsa-mir-199b* | -2.03 | 4.51E-19 |
| *hsa-mir-450a-2* | -1.64 | 4.89E-19 |
| *hsa-mir-511* | -1.30 | 2.74E-18 |
| *hsa-mir-214* | -2.00 | 5.02E-18 |
| *hsa-mir-136* | -1.61 | 1.19E-17 |
| *hsa-mir-486-2* | -1.41 | 1.46E-17 |
| *hsa-mir-486-1* | -1.39 | 4.17E-17 |
| *hsa-mir-411* | -1.54 | 3.98E-16 |
| *hsa-mir-376c* | -1.58 | 2.17E-15 |
| *hsa-mir-369* | -1.60 | 5.62E-15 |
| *hsa-mir-142* | -1.36 | 3.03E-14 |
| *hsa-mir-145* | -1.21 | 3.45E-14 |
| *hsa-mir-223* | -1.11 | 1.58E-13 |
| *hsa-mir-758* | -1.35 | 1.18E-12 |
| *hsa-mir-4686* | -3.43 | 1.71E-12 |
| *hsa-mir-187* | -1.86 | 2.59E-12 |
| *hsa-mir-10a* | -1.15 | 8.20E-12 |
| *hsa-mir-101-2* | -0.87 | 1.30E-11 |
| *hsa-mir-101-1* | -0.87 | 1.36E-11 |
| *hsa-mir-337* | -1.20 | 1.51E-11 |
| *hsa-mir-1247* | -1.62 | 1.92E-11 |
| *hsa-mir-154* | -1.33 | 5.41E-11 |
| *hsa-mir-381* | -1.07 | 6.80E-11 |
| *hsa-mir-654* | -1.25 | 7.04E-11 |
| *hsa-mir-379* | -1.14 | 7.04E-11 |
| *hsa-mir-378c* | -1.07 | 1.07E-10 |
| *hsa-mir-200b* | -1.46 | 1.87E-10 |
| *hsa-mir-6718* | -1.65 | 3.08E-10 |
| *hsa-mir-429* | -1.58 | 3.84E-10 |
| *hsa-mir-200a* | -1.45 | 3.85E-10 |
| *hsa-mir-125b-1* | -1.05 | 4.01E-10 |
| *hsa-mir-299* | -1.19 | 9.18E-10 |
| *hsa-mir-99a* | -1.14 | 1.09E-09 |
| *hsa-mir-125b-2* | -1.03 | 1.25E-09 |
| *hsa-mir-30c-2* | -0.65 | 1.78E-09 |
| *hsa-mir-3614* | -1.11 | 1.87E-09 |
| *hsa-mir-378d-2* | -1.33 | 3.26E-09 |
| *hsa-mir-376a-2* | -1.57 | 5.39E-09 |
| *hsa-let-7c* | -1.08 | 6.87E-09 |
| *hsa-mir-323a* | -1.33 | 1.55E-08 |
| *hsa-mir-134* | -1.04 | 1.62E-08 |
| *hsa-mir-665* | -1.69 | 3.14E-08 |
| *hsa-mir-376a-1* | -1.29 | 3.33E-08 |
| *hsa-mir-150* | -1.14 | 3.86E-08 |
| *hsa-mir-30c-1* | -0.57 | 5.97E-08 |
| *hsa-mir-3653* | -0.99 | 6.68E-08 |
| *hsa-mir-5589* | -1.34 | 6.68E-08 |
| *hsa-mir-30a* | -0.79 | 6.92E-08 |
| *hsa-mir-376b* | -1.14 | 8.38E-08 |
| *hsa-mir-6503* | -1.62 | 9.05E-08 |
| *hsa-mir-627* | -0.87 | 9.71E-08 |
| *hsa-mir-592* | -1.27 | 9.83E-08 |
| *hsa-mir-377* | -1.17 | 1.29E-07 |
| *hsa-mir-378a* | -0.85 | 1.64E-07 |
| *hsa-mir-4772* | -0.96 | 3.63E-07 |
| *hsa-mir-195* | -0.93 | 5.59E-07 |
| *hsa-mir-497* | -0.90 | 6.98E-07 |
| *hsa-mir-874* | -0.87 | 1.76E-06 |
| *hsa-mir-6502* | -1.61 | 2.61E-06 |
| *hsa-mir-621* | -1.78 | 3.59E-06 |
| *hsa-mir-889* | -0.86 | 3.75E-06 |
| *hsa-mir-494* | -1.08 | 4.71E-06 |
| *hsa-mir-33b* | -1.03 | 5.15E-06 |
| *hsa-mir-375* | -0.96 | 5.55E-06 |
| *hsa-mir-4732* | -1.43 | 5.64E-06 |
| *hsa-mir-125a* | -0.74 | 5.74E-06 |
| *hsa-mir-370* | -0.80 | 7.95E-06 |
| *hsa-mir-487b* | -0.84 | 1.34E-05 |
| *hsa-mir-29c* | -0.68 | 1.35E-05 |
| *hsa-mir-505* | -0.68 | 1.62E-05 |
| *hsa-mir-1271* | -0.79 | 2.21E-05 |
| *hsa-mir-503* | -0.66 | 2.99E-05 |
| *hsa-mir-495* | -0.79 | 3.30E-05 |
| *hsa-mir-3607* | -1.19 | 5.06E-05 |
| *hsa-mir-455* | -0.75 | 7.88E-05 |
| *hsa-mir-203a* | -0.78 | 9.02E-05 |
| *hsa-mir-4800* | -1.47 | 1.05E-04 |
| *hsa-mir-655* | -0.88 | 1.12E-04 |
| *hsa-mir-26b* | -0.50 | 1.15E-04 |
| *hsa-mir-7702* | -0.93 | 1.44E-04 |
| *hsa-mir-382* | -0.70 | 1.81E-04 |
| *hsa-mir-543* | -0.82 | 2.37E-04 |
| *hsa-mir-4668* | -0.63 | 2.46E-04 |
| *hsa-mir-127* | -0.65 | 2.69E-04 |
| *hsa-mir-135b* | -0.98 | 4.44E-04 |
| *hsa-mir-141* | -0.90 | 0.00106 |
| *hsa-mir-326* | -0.55 | 0.00107 |
| *hsa-mir-4777* | -1.11 | 0.00124 |
| *hsa-mir-378d-1* | -0.82 | 0.00214 |
| *hsa-mir-133b* | -0.89 | 0.00352 |
| *hsa-mir-23c* | -1.40 | 0.00371 |
| *hsa-mir-485* | -0.61 | 0.00470 |
| *hsa-mir-380* | -0.70 | 0.00537 |
| *hsa-mir-1248* | -0.66 | 0.00826 |
| *hsa-mir-1295b* | -0.54 | 0.00930 |
| *hsa-mir-146a* | -0.50 | 0.01090 |
| *hsa-mir-487a* | -0.56 | 0.01433 |
| *hsa-mir-656* | -0.68 | 0.01647 |
| *hsa-mir-551a* | -1.34 | 0.01679 |
| *hsa-mir-4683* | -1.50 | 0.02946 |
| *hsa-mir-4521* | -0.85 | 0.03027 |
| *hsa-mir-329-2* | -0.80 | 0.03148 |
| *hsa-mir-138-1* | -1.24 | 0.03499 |
| *hsa-mir-4433b* | -0.83 | 0.04241 |
| *hsa-mir-10b* | 4.02 | 1.37E-53 |
| *hsa-mir-3677* | 2.05 | 1.03E-36 |
| *hsa-mir-1269a* | 5.65 | 1.61E-35 |
| *hsa-mir-532* | 1.05 | 2.98E-35 |
| *hsa-mir-589* | 1.33 | 3.62E-33 |
| *hsa-mir-151a* | 1.14 | 1.58E-30 |
| *hsa-mir-106b* | 0.87 | 2.76E-29 |
| *hsa-mir-4746* | 2.25 | 1.86E-25 |
| *hsa-mir-423* | 0.61 | 1.20E-24 |
| *hsa-mir-769* | 0.85 | 1.52E-23 |
| *hsa-mir-93* | 1.13 | 2.39E-23 |
| *hsa-mir-183* | 3.20 | 6.63E-23 |
| *hsa-mir-21* | 1.39 | 9.47E-23 |
| *hsa-mir-501* | 0.98 | 4.88E-22 |
| *hsa-mir-1180* | 1.68 | 9.91E-22 |
| *hsa-mir-500a* | 1.11 | 1.00E-21 |
| *hsa-mir-502* | 0.81 | 3.02E-20 |
| *hsa-mir-221* | 1.29 | 1.70E-19 |
| *hsa-mir-421* | 1.56 | 9.24E-19 |
| *hsa-mir-7706* | 1.61 | 1.28E-17 |
| *hsa-mir-25* | 0.97 | 2.64E-16 |
| *hsa-mir-452* | 2.13 | 3.21E-16 |
| *hsa-mir-5187* | 1.97 | 3.41E-16 |
| *hsa-mir-891a* | 5.07 | 1.12E-15 |
| *hsa-mir-1266* | 2.10 | 8.74E-15 |
| *hsa-mir-182* | 2.66 | 8.75E-15 |
| *hsa-mir-224* | 2.34 | 1.23E-14 |
| *hsa-mir-148b* | 0.73 | 4.39E-14 |
| *hsa-mir-140* | 0.67 | 5.71E-14 |
| *hsa-mir-760* | 2.15 | 5.71E-14 |
| *hsa-mir-4661* | 1.73 | 1.58E-13 |
| *hsa-mir-877* | 1.46 | 4.09E-13 |
| *hsa-mir-96* | 2.79 | 7.04E-13 |
| *hsa-mir-128-1* | 0.86 | 7.69E-13 |
| *hsa-mir-361* | 0.59 | 1.26E-12 |
| *hsa-mir-500b* | 0.75 | 1.30E-12 |
| *hsa-mir-34c* | 2.35 | 1.71E-12 |
| *hsa-mir-1301* | 1.16 | 4.53E-12 |
| *hsa-mir-330* | 1.01 | 5.00E-12 |
| *hsa-mir-222* | 1.10 | 8.20E-12 |
| *hsa-mir-508* | 2.22 | 1.38E-11 |
| *hsa-mir-128-2* | 0.90 | 1.51E-11 |
| *hsa-mir-3682* | 1.59 | 3.21E-11 |
| *hsa-mir-3591* | 2.04 | 4.99E-11 |
| *hsa-mir-660* | 0.64 | 1.62E-10 |
| *hsa-mir-514a-3* | 2.67 | 1.73E-10 |
| *hsa-mir-34a* | 1.12 | 4.57E-10 |
| *hsa-mir-324* | 0.62 | 5.88E-10 |
| *hsa-mir-1307* | 0.83 | 6.53E-10 |
| *hsa-mir-30d* | 0.81 | 2.03E-09 |
| *hsa-mir-188* | 0.88 | 2.07E-09 |
| *hsa-mir-6516* | 1.97 | 4.97E-09 |
| *hsa-mir-671* | 0.71 | 5.39E-09 |
| *hsa-mir-425* | 0.73 | 7.29E-09 |
| *hsa-mir-103a-1* | 0.70 | 8.77E-09 |
| *hsa-mir-4677* | 0.63 | 8.85E-09 |
| *hsa-mir-103a-2* | 0.70 | 9.68E-09 |
| *hsa-mir-514a-1* | 2.55 | 1.97E-08 |
| *hsa-mir-939* | 1.47 | 2.03E-08 |
| *hsa-mir-135a-1* | 2.71 | 2.10E-08 |
| *hsa-mir-185* | 0.58 | 2.39E-08 |
| *hsa-mir-9-3* | 2.57 | 3.52E-08 |
| *hsa-mir-9-1* | 2.58 | 3.63E-08 |
| *hsa-mir-514a-2* | 2.34 | 4.61E-08 |
| *hsa-mir-9-2* | 2.56 | 5.95E-08 |
| *hsa-mir-190b* | 2.45 | 6.68E-08 |
| *hsa-mir-4326* | 0.88 | 9.83E-08 |
| *hsa-mir-1269b* | 4.59 | 1.08E-07 |
| *hsa-mir-3200* | 1.46 | 1.79E-07 |
| *hsa-mir-581* | 1.55 | 2.37E-07 |
| *hsa-mir-937* | 1.46 | 2.58E-07 |
| *hsa-mir-765* | 2.12 | 3.13E-07 |
| *hsa-mir-484* | 0.52 | 3.71E-07 |
| *hsa-mir-5003* | 2.10 | 3.74E-07 |
| *hsa-mir-3928* | 1.20 | 4.30E-07 |
| *hsa-mir-3074* | 0.88 | 4.77E-07 |
| *hsa-mir-107* | 0.51 | 6.03E-07 |
| *hsa-mir-3127* | 1.06 | 6.98E-07 |
| *hsa-mir-3615* | 0.74 | 8.05E-07 |
| *hsa-mir-301b* | 1.51 | 8.51E-07 |
| *hsa-mir-365a* | 0.60 | 9.31E-07 |
| *hsa-mir-3909* | 1.03 | 9.92E-07 |
| *hsa-mir-509-3* | 2.11 | 1.01E-06 |
| *hsa-mir-365b* | 0.60 | 1.18E-06 |
| *hsa-mir-98* | 0.59 | 1.48E-06 |
| *hsa-mir-1304* | 1.27 | 1.54E-06 |
| *hsa-mir-135a-2* | 3.28 | 1.92E-06 |
| *hsa-mir-362* | 0.59 | 2.21E-06 |
| *hsa-mir-132* | 0.66 | 2.65E-06 |
| *hsa-mir-940* | 1.09 | 4.05E-06 |
| *hsa-mir-4664* | 1.80 | 4.80E-06 |
| *hsa-mir-184* | 2.37 | 5.38E-06 |
| *hsa-mir-6755* | 1.02 | 5.98E-06 |
| *hsa-mir-1292* | 1.52 | 1.12E-05 |
| *hsa-mir-550a-1* | 0.80 | 1.15E-05 |
| *hsa-mir-550a-2* | 0.77 | 1.16E-05 |
| *hsa-mir-130b* | 0.89 | 1.97E-05 |
| *hsa-mir-942* | 0.69 | 2.01E-05 |
| *hsa-mir-1270* | 1.59 | 2.21E-05 |
| *hsa-mir-1306* | 0.61 | 2.21E-05 |
| *hsa-mir-509-2* | 2.02 | 2.86E-05 |
| *hsa-mir-6788* | 1.20 | 3.09E-05 |
| *hsa-mir-509-1* | 1.83 | 3.62E-05 |
| *hsa-mir-1226* | 1.13 | 4.50E-05 |
| *hsa-mir-320a* | 0.62 | 4.61E-05 |
| *hsa-mir-652* | 0.64 | 4.68E-05 |
| *hsa-mir-217* | 2.86 | 5.10E-05 |
| *hsa-mir-216a* | 2.72 | 5.51E-05 |
| *hsa-mir-15b* | 0.53 | 6.01E-05 |
| *hsa-mir-504* | 1.09 | 7.39E-05 |
| *hsa-mir-767* | 5.98 | 7.47E-05 |
| *hsa-mir-584* | 0.54 | 9.52E-05 |
| *hsa-mir-548o* | 1.14 | 1.15E-04 |
| *hsa-mir-1229* | 1.36 | 1.63E-04 |
| *hsa-mir-3691* | 1.34 | 2.37E-04 |
| *hsa-mir-3942* | 0.94 | 2.55E-04 |
| *hsa-mir-3610* | 1.21 | 2.74E-04 |
| *hsa-mir-3662* | 1.93 | 3.51E-04 |
| *hsa-mir-1251* | 4.32 | 4.74E-04 |
| *hsa-mir-766* | 0.69 | 6.00E-04 |
| *hsa-mir-664a* | 0.54 | 0.00101 |
| *hsa-mir-552* | 3.89 | 0.00115 |
| *hsa-mir-944* | 1.35 | 0.00117 |
| *hsa-mir-216b* | 2.70 | 0.00118 |
| *hsa-mir-1276* | 1.82 | 0.00127 |
| *hsa-mir-1343* | 0.89 | 0.00133 |
| *hsa-mir-320b-2* | 0.69 | 0.00136 |
| *hsa-mir-105-2* | 5.42 | 0.00138 |
| *hsa-mir-522* | 3.06 | 0.00162 |
| *hsa-mir-34b* | 1.88 | 0.00180 |
| *hsa-mir-32* | 0.53 | 0.00196 |
| *hsa-mir-548o-2* | 0.90 | 0.00207 |
| *hsa-mir-4742* | 0.89 | 0.00214 |
| *hsa-mir-95* | 0.64 | 0.00317 |
| *hsa-mir-548k* | 1.16 | 0.00337 |
| *hsa-mir-5698* | 0.95 | 0.00337 |
| *hsa-mir-5010* | 0.76 | 0.00352 |
| *hsa-mir-212* | 0.56 | 0.00352 |
| *hsa-mir-196b* | 1.31 | 0.00386 |
| *hsa-mir-18a* | 0.55 | 0.00421 |
| *hsa-mir-518c* | 1.65 | 0.00430 |
| *hsa-mir-6875* | 1.21 | 0.00449 |
| *hsa-mir-4786* | 0.73 | 0.00495 |
| *hsa-mir-4784* | 1.25 | 0.00525 |
| *hsa-mir-2114* | 1.75 | 0.00599 |
| *hsa-mir-518e* | 1.28 | 0.00638 |
| *hsa-mir-6806* | 0.58 | 0.00638 |
| *hsa-mir-550a-3* | 0.65 | 0.00663 |
| *hsa-mir-548j* | 0.56 | 0.00716 |
| *hsa-mir-6734* | 1.46 | 0.00751 |
| *hsa-mir-5581* | 0.98 | 0.00766 |
| *hsa-mir-4676* | 0.87 | 0.00826 |
| *hsa-mir-6733* | 0.77 | 0.00863 |
| *hsa-mir-4523* | 1.69 | 0.00872 |
| *hsa-mir-541* | 1.17 | 0.00878 |
| *hsa-mir-2277* | 0.65 | 0.00912 |
| *hsa-mir-518a-1* | 1.10 | 0.00917 |
| *hsa-mir-105-1* | 4.59 | 0.01055 |
| *hsa-mir-219b* | 1.05 | 0.01128 |
| *hsa-mir-6820* | 0.64 | 0.01134 |
| *hsa-mir-643* | 0.76 | 0.01379 |
| *hsa-mir-548s* | 0.90 | 0.01386 |
| *hsa-mir-3940* | 0.55 | 0.01426 |
| *hsa-mir-520a* | 3.96 | 0.01441 |
| *hsa-mir-3912* | 0.52 | 0.01672 |
| *hsa-mir-1288* | 0.72 | 0.01785 |
| *hsa-mir-580* | 0.53 | 0.01900 |
| *hsa-mir-6798* | 1.11 | 0.01974 |
| *hsa-mir-4674* | 1.25 | 0.02481 |
| *hsa-mir-320c-1* | 0.73 | 0.02589 |
| *hsa-mir-5687* | 0.76 | 0.02934 |
| *hsa-mir-4739* | 1.20 | 0.03027 |
| *hsa-mir-5586* | 0.66 | 0.03411 |
| *hsa-mir-7705* | 0.69 | 0.03604 |
| *hsa-mir-4660* | 0.68 | 0.04220 |
| *hsa-mir-4796* | 0.59 | 0.04328 |
| *hsa-mir-527* | 1.08 | 0.04388 |
| *hsa-mir-525* | 1.16 | 0.04415 |
| *hsa-mir-3144* | 2.63 | 0.04470 |
| *hsa-mir-3191* | 0.77 | 0.04763 |
| *hsa-mir-6802* | 0.66 | 0.04870 |
| *hsa-mir-215* | 0.61 | 0.04870 |
| *hsa-mir-4461* | 0.57 | 0.04935 |
| *hsa-mir-6777* | 0.83 | 0.04952 |

**Supplementary Table 4.** 82 pre-miRNAs are significantly dysregulated in zebrafish ABC-HCC by RNA sequencing analysis

| **gene_name** | **log2FoldChange** | **padj** |
| --- | --- | --- |
| *dre-mir-216b* | -5.0748114 | 7.123E-18 |
| *dre-mir-217* | -4.6582354 | 3.4875E-15 |
| *dre-mir-10a* | -4.0313774 | 2.6587E-08 |
| *dre-mir-216a* | -3.7871446 | 1.6051E-11 |
| *dre-mir-206-1* | -2.785726 | 0.00205114 |
| *dre-mir-1-2* | -2.7277176 | 3.476E-05 |
| *dre-mir-107b* | -2.3853405 | 1.2303E-13 |
| *mir196d* | -2.3117162 | 0.04182231 |
| *mir196b* | -2.0327549 | 0.03585138 |
| *dre-mir-181a-2* | -1.9436324 | 7.6676E-12 |
| *dre-mir-459* | -1.7604158 | 0.04618146 |
| *dre-mir-125c* | -1.7237811 | 9.3003E-22 |
| *dre-mir-187-1* | -1.7055529 | 0.00054445 |
| *dre-mir-122* | -1.6956223 | 5.7845E-14 |
| *dre-mir-29b-1* | -1.6792724 | 5.6663E-05 |
| *dre-mir-150* | -1.6419289 | 4.9443E-09 |
| *dre-mir-2188* | -1.4909603 | 0.00013014 |
| *dre-mir-99-1* | -1.299466 | 2.5015E-10 |
| *dre-mir-338-1* | -1.2591732 | 0.0171807 |
| *mir100-2* | -1.2206239 | 5.7013E-05 |
| *dre-mir-730* | -1.2044601 | 0.03585138 |
| *dre-mir-10c* | -1.0985657 | 1.21E-05 |
| *CU442763.1* | -1.0929841 | 0.0217126 |
| *dre-mir-18c* | -1.0739065 | 0.01099649 |
| *dre-mir-7148* | -1.0695306 | 0.00136542 |
| *dre-mir-20b* | -1.0373674 | 0.0052155 |
| *mir99-2* | -1.0235883 | 0.00013129 |
| *dre-mir-181b-2* | -1.01282 | 0.03275326 |
| *dre-mir-22a* | -1.0120652 | 0.00241888 |
| *dre-mir-22b* | -1.0056466 | 1.21E-05 |
| *dre-mir-1388* | -0.9634952 | 0.00193972 |
| *dre-let-7e* | -0.9372929 | 0.00471247 |
| *dre-mir-130a* | -0.9099823 | 0.00013014 |
| *BX088707.1* | -0.7630436 | 0.03585138 |
| *dre-mir-722* | -0.730558 | 0.04484746 |
| *dre-let-7a-2* | -0.6755481 | 0.00867864 |
| *dre-let-7a-6* | -0.6671 | 0.0467683 |
| *CR354430.2* | -0.6177208 | 0.03275326 |
| *dre-mir-107a* | -0.5836311 | 0.03509437 |
| *dre-mir-338-2* | 0.53397522 | 0.04997851 |
| *dre-mir-7b* | 0.70012976 | 0.03509437 |
| *CU550700.1* | 0.73896896 | 0.04182231 |
| *CABZ01078244.1* | 0.75242956 | 0.00426139 |
| *dre-mir-17a-2* | 0.83098844 | 4.689E-06 |
| *dre-mir-429a* | 0.88847823 | 0.00274034 |
| *dre-mir-29a* | 0.93761608 | 0.04293528 |
| *dre-mir-27c-1* | 0.95506741 | 0.01299991 |
| *CABZ01078244.3* | 1.01103918 | 9.5394E-07 |
| *mir24-2* | 1.06403881 | 0.04182231 |
| *dre-mir-93* | 1.21219627 | 1.5802E-06 |
| *dre-mir-25* | 1.22170648 | 0.00205114 |
| *dre-mir-200b* | 1.23153309 | 8.6722E-05 |
| *dre-mir-19d* | 1.25293097 | 1.053E-07 |
| *dre-mir-23a-2* | 1.33498521 | 0.00610725 |
| *BX664751.1* | 1.35882411 | 0.01099221 |
| *dre-mir-16a* | 1.37240093 | 0.00030709 |
| *dre-mir-16b* | 1.39492319 | 1.3169E-05 |
| *dre-mir-200a* | 1.48888876 | 7.3341E-09 |
| *dre-mir-15c* | 1.51468014 | 4.9443E-09 |
| *dre-mir-101a* | 1.53386261 | 8.8228E-06 |
| *dre-mir-15a-1* | 1.57443056 | 0.0007032 |
| *dre-mir-457b* | 1.590706 | 6.2751E-05 |
| *dre-mir-146b* | 1.70878694 | 5.4207E-09 |
| *dre-mir-21-2* | 1.79535044 | 8.734E-07 |
| *dre-mir-146a* | 1.88511823 | 1.2092E-05 |
| *dre-mir-135b* | 1.90939142 | 0.00447049 |
| *dre-mir-457a* | 1.97550534 | 1.4248E-14 |
| *dre-mir-454b* | 2.02624663 | 0.00069887 |
| *dre-mir-21-1* | 2.02965243 | 1.5308E-14 |
| *dre-mir-194b* | 2.07104297 | 0.00022951 |
| *CU550700.2* | 2.22104118 | 0.04182231 |
| *dre-mir-221* | 2.25992984 | 1.0317E-17 |
| *dre-mir-153a* | 2.27188717 | 0.03798538 |
| *dre-mir-2191* | 2.31230544 | 0.01138767 |
| *dre-mir-301b* | 2.37700086 | 0.00088312 |
| *CABZ01078244.2* | 2.48626764 | 8.6608E-22 |
| *dre-mir-222a* | 2.49325799 | 9.5492E-58 |
| *dre-mir-34b* | 2.51418323 | 0.02210025 |
| *dre-mir-15b* | 2.57291167 | 5.2415E-22 |
| *dre-mir-34c* | 2.93502415 | 0.00045007 |
| *dre-mir-130c-2* | 3.74442874 | 1.8194E-05 |
| *dre-mir-190a* | 4.29759093 | 2.6232E-17 |

**Supplementary Table 5**. Homologous miRNAs between human and zebrafish

| **miRNA** | **MASH**  **-HCC** | **ZF** |
| --- | --- | --- |
| *let-7b* | Sig | Sig |
| *let-7e* | Sig | Sig |
| *miR-100* | Sig | Sig |
| *miR-107* | Sig | Sig |
| *miR-10a* | Sig | Sig |
| *miR-146b* | Sig | Sig |
| *miR-15a* | Sig | Sig |
| *miR-15b* | Sig | Sig |
| *miR-194* | Sig | Sig |
| *miR-200b* | Sig | Sig |
| *miR-21* | Sig | Sig |
| *miR-222* | Sig | Sig |
| *miR-23a* | Sig | Sig |
| *miR-24* | Sig | Sig |
| *miR-454* | Sig | Sig |
| *miR-99b* | Sig | Sig |
| *miR-132* | Sig | NS |
| *miR-193b* | Sig | NS |
| *miR-30e* | Sig | NS |
| *miR-1* | NS | Sig |
| *miR-101* | NS | Sig |
| *miR-122* | NS | Sig |
| *miR-130a* | NS | Sig |
| *miR-135b* | NS | Sig |
| *miR-146a* | NS | Sig |
| *miR-150* | NS | Sig |
| *miR-16* | NS | Sig |
| *miR-181a-2* | NS | Sig |
| *miR-181b-2* | NS | Sig |
| *miR-187* | NS | Sig |
| *miR-190a* | NS | Sig |
| *miR-196b* | NS | Sig |
| *miR-200a* | NS | Sig |
| *miR-206* | NS | Sig |
| *miR-20b* | NS | Sig |
| *miR-216a* | NS | Sig |
| *miR-216b* | NS | Sig |
| *miR-217* | NS | Sig |
| *miR-22* | NS | Sig |
| *miR-221* | NS | Sig |
| *miR-25* | NS | Sig |
| *miR-29a* | NS | Sig |
| *miR-29b* | NS | Sig |
| *miR-301b* | NS | Sig |
| *miR-338* | NS | Sig |
| *miR-34b* | NS | Sig |
| *miR-34c* | NS | Sig |
| *miR-429* | NS | Sig |
| *let-7a* | NS | NS |
| *let-7c* | NS | NS |
| *let-7g* | NS | NS |
| *let-7i* | NS | NS |
| *miR-103a* | NS | NS |
| *miR-10b* | NS | NS |
| *miR-125a* | NS | NS |
| *miR-125b* | NS | NS |
| *miR-128-1* | NS | NS |
| *miR-128-2* | NS | NS |
| *miR-130b* | NS | NS |
| *miR-135a* | NS | NS |
| *miR-138* | NS | NS |
| *miR-140* | NS | NS |
| *miR-141* | NS | NS |
| *miR-142* | NS | NS |
| *miR-143* | NS | NS |
| *miR-152* | NS | NS |
| *miR-155* | NS | NS |
| *mIR-17* | NS | NS |
| *miR-181a* | NS | NS |
| *miR-181b* | NS | NS |
| *miR-181c* | NS | NS |
| *miR-182* | NS | NS |
| *miR-183* | NS | NS |
| *miR-18b* | NS | NS |
| *miR-192* | NS | NS |
| *miR-196a* | NS | NS |
| *miR-200c* | NS | NS |
| *miR-202* | NS | NS |
| *miR-203a* | NS | NS |
| *miR-204* | NS | NS |
| *miR-205* | NS | NS |
| *miR-210* | NS | NS |
| *miR-223* | NS | NS |
| *miR-26a* | NS | NS |
| *miR-26b* | NS | NS |
| *miR-27a* | NS | NS |
| *miR-27b* | NS | NS |
| *miR-301a* | NS | NS |
| *miR-30a* | NS | NS |
| *miR-30b* | NS | NS |
| *miR-30d* | NS | NS |
| *miR-31* | NS | NS |
| *miR-363* | NS | NS |
| *miR-375* | NS | NS |
| *miR-455* | NS | NS |
| *miR-489* | NS | NS |
| *miR-9* | NS | NS |
| *miR-92a-1* | NS | NS |
| *miR-93* | NS | NS |
| *miR-96* | NS | NS |

**Supplementary Table 6.** Dysregulated mRNAs in miR-21OE and/or high cholesterol diet conditions.

**Supplementary Table 6a.** WT HCD v WT NCD

| **WT HCD v WT NCD** | | |
| --- | --- | --- |
| **Gene Name** | **L2FC** | **padj** |
| *hmgcra* | -4.3922 | 4.94E-53 |
| *fads2* | -3.4260 | 6.02E-31 |
| *ASS1* | -3.3771 | 1.29E-21 |
| *elovl2* | -2.8117 | 3.91E-20 |
| *cyp51** | -2.8061 | 1.12E-15 |
| *idh1* | -1.4945 | 2.06E-09 |
| *atic* | -1.6490 | 7.86E-09 |
| *abcg1** | 2.4040 | 3.14E-08 |
| *mpx* | 2.4960 | 4.00E-08 |
| *relt* | 1.9234 | 7.20E-08 |
| *chl1a* | 1.8915 | 7.27E-08 |
| *sigmar1* | -1.7397 | 2.20E-07 |
| *abcg5** | 1.5601 | 3.15E-07 |
| *paics* | -1.2866 | 1.45E-06 |
| *nlgn4xb* | -2.1658 | 2.09E-06 |
| *c4b* | 1.6130 | 6.07E-06 |
| *si:ch73-21k16.1* | -2.2141 | 7.22E-06 |
| *mlxipl* | -1.2574 | 7.22E-06 |
| *eef1a1a* | -1.6028 | 7.35E-06 |
| *ldlra* | -1.3629 | 7.48E-06 |
| *fth1b* | -1.5953 | 8.32E-06 |
| *BX539325.2* | 1.6347 | 1.25E-05 |
| *rdh12* | -2.0002 | 1.48E-05 |
| *aclya* | -2.0445 | 1.48E-05 |
| *BX469912.6* | 1.7936 | 1.88E-05 |
| *hsd17b7* | -1.8058 | 1.99E-05 |
| *MCOLN3* | -1.4168 | 1.99E-05 |
| *DOCK4* | -1.9678 | 4.03E-05 |
| *itga2.1* | 2.3554 | 4.88E-05 |
| *zgc:66313* | -2.0981 | 8.28E-05 |
| *elovl5* | -1.5638 | 9.02E-05 |
| *ins* | -0.9980 | 9.02E-05 |
| *sc5d* | -1.5243 | 0.000159 |
| *saa* | 0.1961 | 0.000159 |
| *si:ch211-9d9.1* | 2.1134 | 0.000175 |
| *glud1a* | 1.0839 | 0.000189 |
| *pmt* | -1.6103 | 0.000189 |
| *npas1* | -2.3055 | 0.000199 |
| *lect2l* | 1.5006 | 0.000216 |
| *abcf2b* | -1.3328 | 0.000223 |
| *sele* | 1.6893 | 0.000244 |
| *si:ch211-153b23.5* | 1.8480 | 0.000258 |
| *BX005064.1* | -1.7841 | 0.000336 |
| *itga2.3* | 2.0880 | 0.000339 |
| *zgc:92040* | -0.9441 | 0.000555 |
| *wsb1* | -0.8449 | 0.000555 |
| *agps* | -1.0814 | 0.000555 |
| *tbc1d14* | 1.3162 | 0.000572 |
| *stab2* | 1.8186 | 0.000572 |
| *Lss** | -1.5619 | 0.000572 |
| *NPC1L1** | -1.0615 | 0.000628 |
| *kcnh6b* | 1.5996 | 0.000650 |
| *BX571955.2* | 2.0351 | 0.000683 |
| *GNPAT* | -1.3357 | 0.000692 |
| *ngef* | -0.9080 | 0.001014 |
| *dhcr24* | -1.3969 | 0.001101 |
| *stard4* | -1.1632 | 0.001357 |
| *si:cabz01040626.2* | 0.2096 | 0.001406 |
| *zgc:77938* | -0.8800 | 0.001764 |
| *cntn1a* | 1.1756 | 0.001776 |
| *mmp9* | 1.7437 | 0.001836 |
| *f8* | 1.6547 | 0.001836 |
| *msmo1** | -1.2127 | 0.002001 |
| *fdps* | -1.2987 | 0.002205 |
| *srebf2** | -1.4947 | 0.002205 |
| *zgc:194887* | -1.5146 | 0.002205 |
| *nampta* | -0.8684 | 0.002225 |
| *acod1* | 1.6620 | 0.002384 |
| *swap70b* | 1.5329 | 0.002421 |
| *pltp* | 1.5456 | 0.002426 |
| *HECW1* | -1.1885 | 0.002624 |
| *aacs* | -1.2625 | 0.002642 |
| *abhd15a* | -0.9483 | 0.002642 |
| *lgals9l1* | 1.7718 | 0.002683 |
| *ppm1nb* | -1.8306 | 0.002683 |
| *grasp* | 1.1954 | 0.002886 |
| *casp6b.2* | -0.1569 | 0.002904 |
| *Mvda*fd* | -1.3549 | 0.003134 |
| *acsl1b* | 1.4013 | 0.003501 |
| *fuca1.2* | 1.1531 | 0.003576 |
| *CABZ01079281.1* | 1.7201 | 0.004164 |
| *tmem97* | -1.6684 | 0.004357 |
| *tmem59l* | 1.4249 | 0.004425 |
| *ERG28* | -1.6578 | 0.004425 |
| *acot11b* | -1.2675 | 0.004500 |
| *bcl3* | 1.6945 | 0.004796 |
| *tlcd4a* | -0.8759 | 0.004796 |
| *usp2b* | -1.0451 | 0.004815 |
| *soat2*npc1* | 1.4353 | 0.005680 |
| *nsdhl* | -1.5967 | 0.005747 |
| *mknk2a* | -1.0105 | 0.006026 |
| *sgpl1* | 0.9432 | 0.006147 |
| *cadpsb* | -1.3192 | 0.006147 |
| *CDPF1* | -1.1631 | 0.006288 |
| *zgc:114181* | -0.0321 | 0.0066269 |
| *mat2aa* | -0.9524 | 0.006657 |
| *apoa4a** | 0.8722 | 0.007548 |
| *p2ry13* | 1.4267 | 0.007602 |
| *acss2l* | -1.4806 | 0.007653 |
| *ptp4a3a* | -0.7062 | 0.007813 |
| *tm7sf2* | -1.4266 | 0.007862 |
| *hpxb* | 1.1980 | 0.007862 |
| *nipa2* | 0.9487 | 0.007862 |
| *arhgap12b* | 1.1359 | 0.009374 |
| *ptbp1b* | 1.0149 | 0.009542 |
| *ccdc141* | -0.7009 | 0.009542 |
| *myo15aa* | -1.3435 | 0.009731 |
| *scarb2a* | -1.2971 | 0.009748 |
| *pparaa* | -1.0670 | 0.010447 |
| *lepb* | 0.1022 | 0.010591 |
| *fdft1** | -1.2891 | 0.011292 |
| *junbb* | 1.10045 | 0.011884 |
| *CR855311.1* | 1.3016 | 0.011948 |
| *cyba* | 1.4448 | 0.012926 |
| *si:ch211-152c8.2* | -1.2109 | 0.012926 |
| *kremen1* | 1.2543 | 0.012928 |
| *slitrk2* | 1.5509 | 0.013134 |
| *CABZ01071903.1* | -1.1960 | 0.014162 |
| *tnfrsf1a* | 1.2482 | 0.014175 |
| *dio2* | -1.2879 | 0.014448 |
| *raraa* | -0.9270 | 0.015111 |
| *krt18a.1* | 0.8193 | 0.016468 |
| *cyb5r2* | -1.5365 | 0.016865 |
| *stmn2b* | -1.3705 | 0.018167 |
| *pdxp* | -0.9423 | 0.018325 |
| *MFAP4* | 1.4492 | 0.019413 |
| *ahsg1* | -0.9809 | 0.019877 |
| *hcar1-4* | 1.4358 | 0.021238 |
| *nos1* | -1.0450 | 0.021274 |
| *si:dkey-96l17.6* | 1.4315 | 0.021739 |
| *CABZ01064941.1* | 1.4001 | 0.021739 |
| *tmem176l.2* | 1.2086 | 0.022515 |
| *hck* | 1.4180 | 0.023544 |
| *si:ch211-195m9.3* | 1.4867 | 0.023702 |
| *nrxn1b* | -0.9708 | 0.023989 |
| *ebpl* | -1.3391 | 0.025098 |
| *tspan13a* | -1.1951 | 0.025979 |
| *dao.2* | -1.2628 | 0.026316 |
| *necab2* | -1.2527 | 0.026316 |
| *CABZ01072614.1* | -1.3871 | 0.026693 |
| *sptlc2a* | 1.0443 | 0.026748 |
| *fstl3* | 1.5855 | 0.026990 |
| *hmgcs1** | -1.1506 | 0.028750 |
| *sult3st4* | 1.1777 | 0.028971 |
| *ppp1r13l* | 0.8628 | 0.028997 |
| *mmd* | -0.7332 | 0.029480 |
| *abca1b* | 1.3340 | 0.029783 |
| *psph* | -0.8428 | 0.030401 |
| *decr2* | -1.0057 | 0.030401 |
| *dyrk4* | -1.3653 | 0.030401 |
| *mmp13a* | 1.4718 | 0.030401 |
| *nrp1a* | -0.9044 | 0.030854 |
| *il11a* | 1.2652 | 0.031635 |
| *kif1ab* | -0.8767 | 0.033302 |
| *sec16b* | -0.7832 | 0.033302 |
| *idi1* | -1.2214 | 0.033802 |
| *gstr* | -0.8637 | 0.034801 |
| *cyb5b* | -0.8611 | 0.034888 |
| *si:ch211-212g7.6* | -0.8517 | 0.035842 |
| *zgc:152774* | -1.3938 | 0.036535 |
| *moxd1* | 1.3657 | 0.036752 |
| *zgc:171534* | -1.2584 | 0.039032 |
| *zgc:113142* | 1.1826 | 0.040388 |
| *clcn2a* | 1.4489 | 0.040469 |
| *cxcl8a* | 1.1757 | 0.040778 |
| *mtus1b* | -0.8882 | 0.041753 |
| *necab3* | 1.0077 | 0.042123 |
| *ncam2* | -0.8956 | 0.042643 |
| *sst1.1* | -0.4754 | 0.042718 |
| *mvk* | -1.4908 | 0.046672 |
| *nr1h3* | 0.9776 | 0.046672 |
| *cpe* | -1.2984 | 0.047364 |
| *glula* | 1.0791 | 0.047939 |
| *adam8b* | 0.9169 | 0.048006 |
| *CU693379.1* | 0.7492 | 0.048621 |
| *tuba8l2* | -0.5969 | 0.048887 |
| *pla2g3* | -0.9094 | 0.048966 |
| *acaa1* | -0.7877 | 0.049007 |
| *SLC45A4* | -0.9078 | 0.049007 |
| *lgals9l3* | 1.2948 | 0.049269 |
| *si:ch211-173n18.3* | -1.0762 | 0.049388 |
| *elovl6* | -0.8823 | 0.050779 |

**Supplementary Table 6b.** miR-21OE NCD v WT NCD

| **miR-21OE NCD v WT NCD** | | |
| --- | --- | --- |
| **Gene Name** | **L2FC** | **padj** |
| *nrxn1b* | -2.831 | 7.87E-23 |
| *ldhd* | -2.862 | 2.77E-16 |
| *igfbp2a* | -2.137 | 4.24E-16 |
| *mapk8ip3* | 3.148 | 4.24E-16 |
| *slc23a1* | 2.196 | 4.45E-16 |
| *ret* | 2.684 | 1.48E-15 |
| *si:ch211-113d11.5* | -2.051 | 5.57E-13 |
| *zgc:77439* | -2.347 | 9.66E-12 |
| *ccdc141* | -1.376 | 1.96E-11 |
| *glud1a* | 1.654 | 2.05E-11 |
| *mansc1* | 2.662 | 2.24E-11 |
| *adh8a* | -2.495 | 2.35E-11 |
| *actb1* | 2.091 | 3.58E-11 |
| *ebi3* | -2.132 | 1.13E-10 |
| *rogdi* | -2.012 | 1.41E-10 |
| *dre-mir-30e-2* | 1.622 | 1.66E-10 |
| *aqp9b* | -1.862 | 3.04E-10 |
| *slc27a6* | -2.498 | 5.03E-10 |
| *cnpy3* | -1.607 | 7.61E-10 |
| *bhmt* | 1.190 | 9.57E-10 |
| *itih2* | -1.690 | 1.24E-09 |
| *aifm2* | -1.770 | 1.83E-09 |
| *nme3* | 2.405 | 7.77E-09 |
| *ecpas* | 1.247 | 1.24E-08 |
| *pank1a* | -1.742 | 1.64E-08 |
| *grin2db* | 1.763 | 2.07E-08 |
| *ppm1aa* | -1.020 | 2.26E-07 |
| *nnt* | -1.098 | 2.63E-07 |
| *hsd3b7* | -1.628 | 2.83E-07 |
| *dlat* | 1.242 | 4.26E-07 |
| *sox9a* | -2.056 | 1.54E-06 |
| *rgl1* | 1.198 | 1.59E-06 |
| *angptl2b* | -1.566 | 9.90E-06 |
| *abcc6b.1* | -1.675 | 1.16E-05 |
| *zdhhc3b* | 1.197 | 1.26E-05 |
| *CABZ01092156.1* | 1.267 | 1.29E-05 |
| *nme4* | -1.564 | 1.58E-05 |
| *abcc6b.2* | -1.715 | 1.58E-05 |
| *eml3* | 1.326 | 1.87E-05 |
| *stau2* | 1.022 | 1.94E-05 |
| *slc26a3.2* | -1.036 | 1.97E-05 |
| *pdlim4* | 1.454 | 2.31E-05 |
| *igsf3* | 1.110 | 2.50E-05 |
| *dnase1l41* | 1.648 | 3.01E-05 |
| *pros1* | 1.493 | 3.01E-05 |
| *slc1a3a* | -1.248 | 3.23E-05 |
| *itpr1a* | 1.445 | 3.66E-05 |
| *soul3* | -1.506 | 3.94E-05 |
| *ppp1r3cb* | -1.304 | 4.07E-05 |
| *gabarapb* | -1.123 | 4.57E-05 |
| *soul5* | -1.644 | 4.57E-05 |
| *mdh2* | 0.897 | 4.75E-05 |
| *mgst2* | 1.250 | 4.75E-05 |
| *rhag* | -1.225 | 5.65E-05 |
| *rnf145b* | 1.179 | 6.71E-05 |
| *cox8a* | 0.966 | 7.11E-05 |
| *fabp7a* | 1.638 | 7.55E-05 |
| *acot11b* | -1.552 | 8.67E-05 |
| *dusp2* | -1.669 | 1.00E-04 |
| *slc16a6b* | -1.323 | 1.10E-04 |
| *agmo* | 1.161 | 1.10E-04 |
| *abhd15a* | -1.095 | 1.17E-04 |
| *nucb2a* | -0.938 | 1.39E-04 |
| *actr2a* | 0.927 | 1.61E-04 |
| *aldh9a1a1* | -1.243 | 1.66E-04 |
| *aadat* | -0.932 | 1.69E-04 |
| *npas1* | -2.286 | 1.85E-04 |
| *dre-mir-21-1* | 1.734 | 2.36E-04 |
| *maats1* | -0.774 | 2.62E-04 |
| *ncl* | 1.498 | 2.62E-04 |
| *tpp1* | 1.020 | 2.80E-04 |
| *agbl1* | 0.999 | 3.00E-04 |
| *snx13* | 0.964 | 3.23E-04 |
| *slc43a2a* | 1.249 | 3.39E-04 |
| *si:ch211-122f10.4* | 0.927 | 3.39E-04 |
| *serpinb1l3* | 1.082 | 3.46E-04 |
| *hvcn1* | -1.133 | 3.85E-04 |
| *tgfbi* | -1.468 | 3.89E-04 |
| *hook2* | 1.196 | 3.91E-04 |
| *GALNT10* | 0.915 | 3.91E-04 |
| *CT583728.24* | -0.186 | 3.91E-04 |
| *hmgcl* | -0.965 | 3.92E-04 |
| *ube2g2* | 1.111 | 4.17E-04 |
| *dab2* | 1.371 | 4.17E-04 |
| *pacs2* | 1.107 | 4.24E-04 |
| *lgals2b* | 0.831 | 4.42E-04 |
| *vdra* | -1.099 | 4.80E-04 |
| *etv4* | 1.697 | 5.46E-04 |
| *pdzrn3a* | 1.260 | 5.54E-04 |
| *cadpsb* | -1.491 | 5.58E-04 |
| *jph1a* | 1.273 | 6.83E-04 |
| *taldo1* | -0.757 | 7.58E-04 |
| *gadd45aa* | -1.359 | 8.58E-04 |
| *l3mbtl3* | -1.039 | 9.50E-04 |
| *atp5if1a* | 1.073 | 1.03E-03 |
| *hgd* | -0.969 | 1.04E-03 |
| *sec31b* | 0.768 | 1.10E-03 |
| *paics* | -0.957 | 1.10E-03 |
| *npr1b* | 1.288 | 1.13E-03 |
| *gba* | 0.971 | 1.17E-03 |
| *rbp5* | 1.481 | 1.24E-03 |
| *PSTK* | 1.113 | 1.30E-03 |
| *hbl4* | -1.100 | 1.34E-03 |
| *got2a* | -0.813 | 1.36E-03 |
| *si:ch211-186e20.2* | -1.306 | 1.36E-03 |
| *atp7b* | -1.226 | 1.40E-03 |
| *pklr* | -1.105 | 1.50E-03 |
| *prkcg* | 1.705 | 1.52E-03 |
| *pdzd3a* | 1.286 | 1.52E-03 |
| *rhobtb2a* | -1.240 | 1.52E-03 |
| *pdk1* | 1.178 | 1.63E-03 |
| *fam120a* | 1.090 | 1.63E-03 |
| *CT027815.1* | -1.053 | 1.63E-03 |
| *lrit3a* | 1.613 | 1.68E-03 |
| *ptdss2* | -1.195 | 1.73E-03 |
| *hmgn6* | 1.080 | 1.74E-03 |
| *lpcat3* | 0.915 | 1.74E-03 |
| *mt2* | 1.564 | 1.76E-03 |
| *si:ch73-364h19.2* | 1.394 | 1.76E-03 |
| *nfil3-6* | -1.372 | 1.97E-03 |
| *aco2* | 0.692 | 2.13E-03 |
| *txndc12* | 0.963 | 2.24E-03 |
| *gnpat* | 1.038 | 2.25E-03 |
| *cyp2x9* | -1.391 | 2.27E-03 |
| *ugt5g1* | 0.824 | 2.48E-03 |
| *n4bp3* | -0.943 | 2.52E-03 |
| *ap2b1* | 0.702 | 2.60E-03 |
| *dhx32a* | 1.369 | 2.60E-03 |
| *VDR* | -1.034 | 2.60E-03 |
| *csad* | -1.243 | 2.76E-03 |
| *slc52a3* | 1.473 | 2.86E-03 |
| *g6pd* | -0.935 | 2.90E-03 |
| *RAMP3* | 0.861 | 2.99E-03 |
| *sts* | 1.164 | 3.15E-03 |
| *slc2a13b* | 0.961 | 3.15E-03 |
| *rhoab* | 1.330 | 3.24E-03 |
| *ggcx* | -0.981 | 3.29E-03 |
| *pdhb* | -0.696 | 3.31E-03 |
| *pcgf5b* | -1.376 | 3.46E-03 |
| *dpp6b* | 1.514 | 3.70E-03 |
| *hibadha* | 1.051 | 4.21E-03 |
| *tsc1b* | 1.087 | 4.21E-03 |
| *si:ch73-269m23.5* | -0.904 | 4.32E-03 |
| *zgc:110410* | 0.985 | 4.55E-03 |
| *hlfa* | -0.713 | 4.69E-03 |
| *frmd4bb* | -1.199 | 4.69E-03 |
| *ahsg2* | -1.397 | 4.86E-03 |
| *hook2* | 1.060 | 4.94E-03 |
| *max* | 0.760 | 4.97E-03 |
| *ppfibp1a* | -0.831 | 4.97E-03 |
| *zgc:194887* | -1.452 | 4.97E-03 |
| *coq8aa* | -0.737 | 5.18E-03 |
| *gart* | -0.962 | 5.23E-03 |
| *zgc:91944* | -0.825 | 5.23E-03 |
| *zgc:110339* | 1.005 | 5.23E-03 |
| *scmh1* | -1.373 | 5.40E-03 |
| *si:dkey-18a10.3* | 0.870 | 5.40E-03 |
| *ndrg3b* | 0.932 | 5.57E-03 |
| *slc38a6* | 1.006 | 5.64E-03 |
| *enosf1* | -1.164 | 5.93E-03 |
| *sod3a* | -1.056 | 5.94E-03 |
| *abhd3* | -1.325 | 6.05E-03 |
| *zgc:153031* | 1.006 | 6.05E-03 |
| *slc16a9a* | 1.477 | 6.10E-03 |
| *ephx2* | -0.802 | 6.63E-03 |
| *ywhaqb* | 0.677 | 6.68E-03 |
| *capza1a* | 0.849 | 6.68E-03 |
| *slc51a* | 0.882 | 6.68E-03 |
| *sgpl1* | 0.908 | 6.68E-03 |
| *ubac2* | -1.056 | 6.68E-03 |
| *ubap2b* | 0.707 | 6.72E-03 |
| *apooa* | -0.955 | 6.72E-03 |
| *slc47a3* | 0.892 | 6.72E-03 |
| *acy3.2* | 1.034 | 6.82E-03 |
| *slc16a13* | -1.159 | 7.44E-03 |
| *rhoaa* | 0.768 | 7.48E-03 |
| *cemip* | -1.595 | 7.61E-03 |
| *mettl9* | 0.815 | 7.64E-03 |
| *gucd1* | 0.926 | 7.64E-03 |
| *sms* | 1.101 | 7.69E-03 |
| *ktn1* | 0.736 | 7.73E-03 |
| *hps1* | -0.984 | 7.92E-03 |
| *bfb* | 1.128 | 7.92E-03 |
| *zgc:66313* | -1.476 | 7.92E-03 |
| *CU693379.1* | 0.845 | 7.92E-03 |
| *aldh9a1a.2* | -1.132 | 8.04E-03 |
| *slc26a4* | -1.628 | 8.04E-03 |
| *f7* | -0.741 | 8.07E-03 |
| *acadm* | -0.823 | 8.07E-03 |
| *dnah1* | -1.459 | 8.08E-03 |
| *si:ch211-133n4.9* | -1.400 | 8.15E-03 |
| *fam214a* | 0.872 | 8.20E-03 |
| *fbxo8* | -1.012 | 8.58E-03 |
| *tmem59* | 0.687 | 8.77E-03 |
| *spag6* | 1.602 | 8.84E-03 |
| *sult3st3* | -1.329 | 8.98E-03 |
| *BX511227.4* | 1.351 | 9.11E-03 |
| *tmem63ba* | 0.652 | 9.11E-03 |
| *laptm4b* | 1.019 | 9.24E-03 |
| *cldn15la* | 1.247 | 9.50E-03 |
| *faah2a* | 0.951 | 9.58E-03 |
| *twf2a* | -1.250 | 9.58E-03 |
| *mlxipl* | -0.840 | 9.58E-03 |
| *cthl* | -1.460 | 9.64E-03 |
| *dhrs13a.1* | -0.655 | 9.85E-03 |
| *raver2* | -1.263 | 9.85E-03 |
| *cyp4v7* | -0.997 | 1.01E-02 |
| *ugt2a7* | 1.021 | 1.03E-02 |
| *CABZ01057057.1* | -0.755 | 1.03E-02 |
| *atp2a2a* | -0.971 | 1.06E-02 |
| *pparab* | -0.979 | 1.13E-02 |
| *acss3* | -0.884 | 1.14E-02 |
| *rbp2b* | 1.050 | 1.14E-02 |
| *igfbp3* | -1.288 | 1.14E-02 |
| *ctsd* | 0.841 | 1.18E-02 |
| *pcxb* | -0.899 | 1.19E-02 |
| *si:dkey-239n17.6* | 1.258 | 1.20E-02 |
| *wu:fj16a03* | 0.696 | 1.20E-02 |
| *syngr2a* | 0.702 | 1.22E-02 |
| *si:rp71-79p20.2* | -1.145 | 1.23E-02 |
| *sec24a* | -0.566 | 1.23E-02 |
| *CR388178.1* | -0.718 | 1.23E-02 |
| *zfp36l1b* | -0.947 | 1.24E-02 |
| *JMJD8* | -1.358 | 1.29E-02 |
| *MCOLN3* | 0.952 | 1.32E-02 |
| *dnajc16l* | 1.012 | 1.32E-02 |
| *CABZ01084501.2* | 0.787 | 1.33E-02 |
| *RASGRP2* | -1.018 | 1.36E-02 |
| *si:dkey-239i20.2* | -1.016 | 1.39E-02 |
| *cryaa* | -0.641 | 1.39E-02 |
| *si:ch1073-280e3.1* | 1.617 | 1.39E-02 |
| *npc2* | -0.876 | 1.39E-02 |
| *sdhc* | 0.787 | 1.41E-02 |
| *zgc:174259* | -1.245 | 1.41E-02 |
| *ek1* | 0.953 | 1.42E-02 |
| *tbc1d10b* | 0.742 | 1.42E-02 |
| *prelid3b* | -0.987 | 1.43E-02 |
| *nqo1* | -0.876 | 1.43E-02 |
| *lepb* | 0.081 | 1.43E-02 |
| *abhd11* | -0.894 | 1.46E-02 |
| *grik1a* | 1.350 | 1.48E-02 |
| *zfx* | -0.950 | 1.48E-02 |
| *wsb1* | -0.651 | 1.52E-02 |
| *cebpa* | -1.128 | 1.54E-02 |
| *acsl1b* | 1.194 | 1.55E-02 |
| *me1* | -0.991 | 1.55E-02 |
| *pgap3* | 0.991 | 1.56E-02 |
| *pank1b* | -0.856 | 1.56E-02 |
| *reep3a* | 0.991 | 1.58E-02 |
| *bbox1* | -0.969 | 1.58E-02 |
| *fhdc5* | -1.231 | 1.62E-02 |
| *sfxn1* | 0.735 | 1.62E-02 |
| *ASS1* | -1.110 | 1.67E-02 |
| *smyhc1* | -1.517 | 1.69E-02 |
| *aldob* | -0.856 | 1.70E-02 |
| *cyp27a7* | -1.031 | 1.71E-02 |
| *cat* | -0.976 | 1.73E-02 |
| *psmd10* | -1.209 | 1.74E-02 |
| *itm2bb* | -0.734 | 1.74E-02 |
| *si:rp71-36a1.1* | -1.235 | 1.76E-02 |
| *zgc:77938* | 0.705 | 1.86E-02 |
| *atp1b1a* | 0.825 | 1.87E-02 |
| *cacna1db* | -0.694 | 1.91E-02 |
| *vps26a* | 1.394 | 1.91E-02 |
| *wdr89* | -0.866 | 1.93E-02 |
| *fam83d* | -1.419 | 1.96E-02 |
| *adkb* | -1.025 | 1.97E-02 |
| *hmgb3a* | 1.246 | 1.98E-02 |
| *fdxr* | 0.815 | 1.98E-02 |
| *EML4* | 0.697 | 1.98E-02 |
| *pak2b* | 0.695 | 2.01E-02 |
| *BX936439.1* | 0.918 | 2.01E-02 |
| *zgc:154142* | 1.212 | 2.02E-02 |
| *BX469912.6* | 0.971 | 2.04E-02 |
| *itih3b* | -0.707 | 2.04E-02 |
| *cratb* | 0.949 | 2.10E-02 |
| *smox* | 0.988 | 2.12E-02 |
| *frem2b* | 1.321 | 2.21E-02 |
| *acox1* | -0.814 | 2.28E-02 |
| *si:ch73-103b11.2* | -0.579 | 2.31E-02 |
| *acot19* | 1.264 | 2.33E-02 |
| *fam120b* | 0.891 | 2.35E-02 |
| *CABZ01072254.1* | -0.808 | 2.40E-02 |
| *mpp6a* | -0.972 | 2.41E-02 |
| *si:ch211-221f10.2* | -0.790 | 2.41E-02 |
| *FP101887.1* | -0.973 | 2.42E-02 |
| *ube2v1* | 0.636 | 2.42E-02 |
| *tmem88a* | -0.714 | 2.46E-02 |
| *lonrf1l* | -0.906 | 2.46E-02 |
| *mif* | -0.872 | 2.48E-02 |
| *ugt5a1* | 0.854 | 2.48E-02 |
| *flrt2* | -0.911 | 2.55E-02 |
| *kcng4a* | 1.334 | 2.68E-02 |
| *pfn2* | 0.617 | 2.79E-02 |
| *serhl* | 0.904 | 2.79E-02 |
| *zgc:174356* | 1.181 | 2.79E-02 |
| *cyp46a1.3* | -1.411 | 2.79E-02 |
| *apoc4* | -1.077 | 2.92E-02 |
| *arl13a* | -1.174 | 3.13E-02 |
| *zgc:103759* | -1.412 | 3.31E-02 |
| *mpp1* | 0.815 | 3.41E-02 |
| *si:ch211-288g17.4* | -0.693 | 3.44E-02 |
| *cemip2* | 0.642 | 3.44E-02 |
| *proca* | -0.626 | 3.47E-02 |
| *rfng* | 0.991 | 3.49E-02 |
| *si:dkey-246g23.4* | 1.022 | 3.49E-02 |
| *hsd11b1lb* | -0.709 | 3.49E-02 |
| *nob1* | -1.214 | 3.49E-02 |
| *zgc:153921* | 0.680 | 3.49E-02 |
| *b3gnt5b* | 1.198 | 3.49E-02 |
| *bnip3lb* | 0.864 | 3.53E-02 |
| *slc40a1* | 1.059 | 3.62E-02 |
| *rapgef3* | -0.970 | 3.62E-02 |
| *si:ch211-153b23.7* | -1.211 | 3.72E-02 |
| *l2hgdh* | 1.136 | 3.73E-02 |
| *dchs1b* | -0.667 | 3.73E-02 |
| *dpy19l1l* | 0.730 | 3.75E-02 |
| *spata20* | -0.677 | 3.85E-02 |
| *ghrhra* | -1.402 | 3.85E-02 |
| *lrrtm1* | -0.761 | 3.85E-02 |
| *si:ch211-160o17.2* | 1.195 | 3.85E-02 |
| *si:dkey-104n9.1* | -1.269 | 3.86E-02 |
| *hsd17b4* | -0.856 | 3.89E-02 |
| *dio1* | -0.820 | 4.00E-02 |
| *xrcc1* | 0.865 | 4.00E-02 |
| *tep1* | 0.663 | 4.10E-02 |
| *prkchb* | -1.217 | 4.13E-02 |
| *sik2b* | -0.772 | 4.18E-02 |
| *oaz2a* | -0.981 | 4.23E-02 |
| *CLSTN2* | -1.004 | 4.27E-02 |
| *gsap* | 1.302 | 4.27E-02 |
| *psen1* | 0.719 | 4.30E-02 |
| *gcshb* | 0.755 | 4.30E-02 |
| *CABZ01060030.2* | -1.278 | 4.30E-02 |
| *arrdc3a* | 1.048 | 4.30E-02 |
| *dnajc18* | -0.748 | 4.30E-02 |
| *fuca1.2* | 0.899 | 4.32E-02 |
| *mmp17a* | 1.062 | 4.32E-02 |
| *poldip2* | -0.632 | 4.32E-02 |
| *ces3* | -1.021 | 4.36E-02 |
| *dgkb* | -1.225 | 4.37E-02 |
| *clptm1* | 0.840 | 4.39E-02 |
| *calm3a* | 0.835 | 4.40E-02 |
| *rnd1b* | -1.142 | 4.49E-02 |
| *txn* | -0.791 | 4.49E-02 |
| *cth* | -0.828 | 4.49E-02 |
| *si:ch211-12h2.8* | -0.787 | 4.53E-02 |
| *anapc7* | 0.994 | 4.53E-02 |
| *kctd13* | 0.860 | 4.59E-02 |
| *fah* | -0.657 | 4.60E-02 |
| *slc47a2.1* | -1.033 | 4.61E-02 |
| *hp* | -0.415 | 4.64E-02 |
| *cyp2k19* | -1.041 | 4.65E-02 |
| *si:ch211-214j24.15* | -0.615 | 4.69E-02 |
| *trappc12* | 0.988 | 4.69E-02 |
| *apool* | -0.656 | 4.69E-02 |
| *plch2a* | 1.416 | 4.72E-02 |
| *vps53* | 0.853 | 4.79E-02 |
| *slc7a9* | -1.108 | 4.82E-02 |
| *arl4aa* | 1.232 | 4.90E-02 |
| *ctsba* | -0.541 | 4.96E-02 |
| *tmem30c* | 0.735 | 5.03E-02 |

**Supplementary Table 6c.** miR-21OE HCD v WT HCD

| **miR-21OE HCD v WT HCD** | | |
| --- | --- | --- |
| **Gene Name** | **L2FC** | **padj** |
| *slc23a1* | 2.323 | 3.15E-18 |
| *BX539325.2* | -2.742 | 2.76E-16 |
| *grin2db* | 2.407 | 5.02E-16 |
| *mansc1* | 3.114 | 6.04E-16 |
| *mapk8ip3* | 2.935 | 9.06E-14 |
| *bhmt* | 1.378 | 4.74E-13 |
| *aifm2* | -2.001 | 3.86E-12 |
| *nrxn1b* | -2.066 | 5.18E-12 |
| *dio2* | 2.721 | 1.61E-11 |
| *ldhd* | -2.304 | 7.78E-11 |
| *adh8a* | -2.419 | 8.93E-11 |
| *angptl2b* | -2.055 | 1.00E-10 |
| *rogdi* | -2.020 | 1.00E-10 |
| *si:ch211-113d11.5* | -1.865 | 1.00E-10 |
| *aqp9b* | -1.856 | 3.69E-10 |
| *actb1* | 1.980 | 7.80E-10 |
| *igfbp2a* | -1.681 | 7.80E-10 |
| *fads2* | 1.942 | 9.81E-10 |
| *CT027815.1* | -1.684 | 1.01E-09 |
| *ret* | 2.100 | 1.70E-09 |
| *slc27a6* | -2.427 | 2.43E-09 |
| *raver2* | -2.070 | 2.52E-07 |
| *hsd3b7* | -1.636 | 2.77E-07 |
| *wu:fj16a03* | 1.130 | 5.07E-07 |
| *nme3* | 2.216 | 5.29E-07 |
| *stau2* | 1.140 | 6.56E-07 |
| *ppm1aa* | -0.986 | 6.57E-07 |
| *MCOLN3* | 1.559 | 6.90E-07 |
| *CR450764.18* | -0.002 | 7.24E-07 |
| *hook2* | 1.505 | 7.86E-07 |
| *eml3* | 1.452 | 9.84E-07 |
| *itih2* | -1.408 | 1.89E-06 |
| *limk1a* | 1.258 | 1.91E-06 |
| *cnpy3* | -1.306 | 2.15E-06 |
| *slc16a6b* | 1.333 | 2.38E-06 |
| *hbl4* | -1.430 | 3.24E-06 |
| *ebi3* | -1.633 | 4.51E-06 |
| *slc25a48* | -1.908 | 4.74E-06 |
| *tmem59l* | 1.998 | 4.84E-06 |
| *rnf145b* | 1.287 | 5.41E-06 |
| *GALNT10* | 1.083 | 6.43E-06 |
| *gabarapb* | -1.195 | 6.88E-06 |
| *igfbp3* | -1.934 | 1.25E-05 |
| *hook2* | 1.419 | 1.32E-05 |
| *RAMP3* | 1.101 | 1.55E-05 |
| *dre-mir-30e-2* | 1.361 | 1.84E-05 |
| *itpr1a* | 1.469 | 2.15E-05 |
| *si:dkey-58b18.9* | -0.025 | 2.15E-05 |
| *fam120a* | 1.335 | 2.30E-05 |
| *CABZ01092156.1* | 1.205 | 3.58E-05 |
| *ube2g2* | 1.232 | 3.96E-05 |
| *si:ch211-153b23.7* | -1.839 | 4.69E-05 |
| *agmo* | -1.371 | 4.79E-05 |
| *glud1a* | 1.128 | 4.81E-05 |
| *mgst2* | 1.227 | 5.41E-05 |
| *dnah1* | -1.917 | 5.41E-05 |
| *p4ha3* | 1.505 | 5.54E-05 |
| *mdh2* | 0.888 | 5.78E-05 |
| *dlat* | 1.038 | 7.32E-05 |
| *zgc:100868* | -1.980 | 7.45E-05 |
| *crybb1l1* | -0.088 | 7.91E-05 |
| *zgc:77439* | -1.546 | 8.00E-05 |
| *pklr* | -1.270 | 8.19E-05 |
| *jph1a* | 1.392 | 8.88E-05 |
| *si:zfos-943e10.1* | -1.214 | 1.07E-04 |
| *abcc6b.1* | -1.519 | 1.40E-04 |
| *pacs2* | 1.150 | 1.41E-04 |
| *tmem63ba* | 0.833 | 1.54E-04 |
| *si:rp71-36a1.1* | -1.673 | 1.90E-04 |
| *mylipb* | 1.704 | 1.90E-04 |
| *apooa* | 1.093 | 2.05E-04 |
| *si:dkey-96l17.6* | -1.842 | 2.29E-04 |
| *zgc:77938* | 0.942 | 2.93E-04 |
| *soul5* | -1.461 | 2.95E-04 |
| *pak2b* | 0.929 | 2.96E-04 |
| *slc1a3a* | -1.123 | 3.34E-04 |
| *txndc12* | 1.066 | 3.49E-04 |
| *ktn1* | 0.884 | 4.63E-04 |
| *alox5b.3* | -1.719 | 4.63E-04 |
| *abcc6b.2* | -1.501 | 4.66E-04 |
| *ywhag1* | 1.350 | 4.90E-04 |
| *pcgf5b* | -1.543 | 5.45E-04 |
| *atp5if1a* | 1.100 | 5.45E-04 |
| *ek1* | 1.188 | 6.42E-04 |
| *arl13a* | -1.574 | 6.57E-04 |
| *nacc1a* | 0.972 | 7.51E-04 |
| *si:ch73-364h19.2* | 1.449 | 7.69E-04 |
| *nnt* | -0.812 | 7.88E-04 |
| *ubap2b* | 0.814 | 7.99E-04 |
| *sec31b* | 0.773 | 9.13E-04 |
| *sult3st3* | -1.553 | 9.13E-04 |
| *si:ch211-67e16.11* | 2.019 | 9.13E-04 |
| *scmh1* | 0.943 | 9.77E-04 |
| *maats1* | 1.373 | 1.42E-03 |
| *pank1a* | -1.156 | 1.50E-03 |
| *atp1a1a.3* | -1.665 | 1.50E-03 |
| *igsf3* | 0.907 | 1.51E-03 |
| *ankrd9* | 1.621 | 1.52E-03 |
| *clockb* | -0.991 | 1.53E-03 |
| *usp2b* | 1.096 | 1.53E-03 |
| *sccpdha.1* | 1.611 | 1.53E-03 |
| *nos1* | 1.220 | 1.55E-03 |
| *rnf175* | 1.246 | 1.56E-03 |
| *si:dkey-16p21.8* | 1.077 | 1.71E-03 |
| *slc25a11* | 0.945 | 1.79E-03 |
| *ecpas* | 0.814 | 1.83E-03 |
| *neto2b* | 1.704 | 1.97E-03 |
| *itga2.3* | -1.806 | 2.02E-03 |
| *laptm4b* | 1.118 | 2.03E-03 |
| *egr2a* | 1.578 | 2.03E-03 |
| *hmgn6* | 1.060 | 2.23E-03 |
| *eps8l3a* | -1.445 | 2.23E-03 |
| *sfxn1* | 0.864 | 2.24E-03 |
| *stmn2b* | 1.550 | 2.53E-03 |
| *snx13* | 0.852 | 2.61E-03 |
| *pdlim4* | 1.117 | 2.84E-03 |
| *pstk* | 1.046 | 2.84E-03 |
| *mthfd2* | 1.125 | 2.90E-03 |
| *prkcg* | 1.644 | 3.12E-03 |
| *acy3.2* | 1.095 | 3.12E-03 |
| *ndrg3b* | 0.975 | 3.12E-03 |
| *si:dkey-6b12.5* | 1.303 | 3.12E-03 |
| *pgbd5* | -1.254 | 3.32E-03 |
| *rbp5* | 1.377 | 3.39E-03 |
| *slc16a9a* | 1.540 | 3.41E-03 |
| *mif* | 1.016 | 3.41E-03 |
| *CR388178.1* | -0.808 | 3.41E-03 |
| *ryr2a* | 1.288 | 3.42E-03 |
| *galnt10* | 1.009 | 3.42E-03 |
| *gphna* | 0.982 | 3.43E-03 |
| *dab2* | 1.216 | 3.57E-03 |
| *dap1b* | 0.766 | 3.57E-03 |
| *mpx* | -1.517 | 3.65E-03 |
| *elovl2* | 1.148 | 3.68E-03 |
| *lipf* | 0.968 | 3.69E-03 |
| *nucb2a* | -0.777 | 3.96E-03 |
| *mospd2* | 0.876 | 4.11E-03 |
| *unm_sa1261* | 1.428 | 4.42E-03 |
| *slc2a5* | -1.492 | 4.50E-03 |
| *pcxb* | -0.976 | 4.63E-03 |
| *tbc1d10b* | 0.810 | 4.70E-03 |
| *bfb* | -1.027 | 4.79E-03 |
| *egf* | 1.384 | 4.79E-03 |
| *fdxr* | 0.925 | 4.79E-03 |
| *txn* | -0.973 | 4.86E-03 |
| *caspa* | -1.447 | 5.06E-03 |
| *si:ch211-28p3.4* | -0.020 | 5.21E-03 |
| *egln3* | 1.659 | 5.43E-03 |
| *cbr1l* | -1.046 | 5.44E-03 |
| *ctnna2* | 1.399 | 5.55E-03 |
| *atg16l1* | 0.761 | 5.58E-03 |
| *CT583728.24* | -0.036 | 5.84E-03 |
| *TENM2* | 1.452 | 5.87E-03 |
| *BX511227.4* | 1.394 | 5.87E-03 |
| *tmem59* | 0.708 | 6.36E-03 |
| *gart* | 0.988 | 6.49E-03 |
| *ggcx* | -0.937 | 6.49E-03 |
| *lamc3* | -1.501 | 6.56E-03 |
| *gcshb* | 0.901 | 6.83E-03 |
| *hsd11b1lb* | 0.790 | 7.43E-03 |
| *ugt5g1* | 0.761 | 7.66E-03 |
| *slc2a13b* | 0.892 | 7.66E-03 |
| *cldn15la* | 1.245 | 7.66E-03 |
| *phtf2* | 0.990 | 8.31E-03 |
| *fosb* | 1.470 | 8.58E-03 |
| *vav3b* | -1.262 | 8.63E-03 |
| *trib1* | 1.088 | 8.77E-03 |
| *acss3* | -0.912 | 8.80E-03 |
| *ctbp2a* | 0.775 | 8.83E-03 |
| *necab3* | -1.128 | 8.83E-03 |
| *ada* | -1.209 | 8.84E-03 |
| *VDR* | -0.948 | 8.96E-03 |
| *adcy3a* | 0.931 | 9.34E-03 |
| *slc13a5a* | -1.138 | 9.34E-03 |
| *si:dkey-83k24.5* | 0.966 | 9.34E-03 |
| *entpd5b* | 0.932 | 9.43E-03 |
| *fgb* | -0.659 | 9.47E-03 |
| *akr1a1a* | 0.993 | 9.47E-03 |
| *zgc:174356* | 1.335 | 9.47E-03 |
| *BX547928.1* | 1.553 | 9.47E-03 |
| *si:ch1073-126c3.2* | -1.123 | 9.54E-03 |
| *bnip3lb* | 0.977 | 9.63E-03 |
| *zgc:153031* | 0.969 | 9.82E-03 |
| *anapc7* | -0.911 | 1.01E-02 |
| *soul3* | -1.092 | 1.01E-02 |
| *lrit3a* | 1.561 | 1.03E-02 |
| *fabp7a* | 1.265 | 1.03E-02 |
| *cyp2k18* | -1.680 | 1.04E-02 |
| *rnd2* | 1.308 | 1.13E-02 |
| *twf2a* | -1.231 | 1.13E-02 |
| *slc12a10.2* | -1.473 | 1.13E-02 |
| *zgc:162183* | 0.987 | 1.14E-02 |
| *cyp4v7* | -0.987 | 1.16E-02 |
| *si:ch211-186e20.2* | -1.127 | 1.19E-02 |
| *hmgcl* | -0.772 | 1.22E-02 |
| *naprt* | 0.775 | 1.22E-02 |
| *KIF2A* | 0.762 | 1.27E-02 |
| *ppm1h* | 1.022 | 1.27E-02 |
| *dpy19l1l* | 0.813 | 1.28E-02 |
| *ipmka* | 1.212 | 1.30E-02 |
| *BX324177.12* | -1.071 | 1.30E-02 |
| *sult3st2* | -1.215 | 1.32E-02 |
| *slc3a2b* | 0.919 | 1.34E-02 |
| *slc25a12* | 0.706 | 1.35E-02 |
| *fbxo8* | -0.972 | 1.36E-02 |
| *zgc:66479* | -1.446 | 1.36E-02 |
| *pgap3* | 1.006 | 1.36E-02 |
| *zdhhc3b* | 0.803 | 1.36E-02 |
| *ppp1r3ca* | -1.493 | 1.36E-02 |
| *EML4* | 0.723 | 1.36E-02 |
| *kremen1* | 1.165 | 1.36E-02 |
| *sdhb* | 0.647 | 1.36E-02 |
| *cemip2* | 0.700 | 1.37E-02 |
| *FO704750.1* | 0.983 | 1.37E-02 |
| *il11ra* | -0.740 | 1.40E-02 |
| *cep170b* | 0.659 | 1.40E-02 |
| *slc38a6* | 0.922 | 1.41E-02 |
| *si:ch211-122f10.4* | 0.704 | 1.42E-02 |
| *agbl1* | 0.761 | 1.50E-02 |
| *tmod4* | 1.098 | 1.51E-02 |
| *itm2bb* | -0.749 | 1.51E-02 |
| *acot11b* | -1.127 | 1.54E-02 |
| *ddt* | -0.715 | 1.58E-02 |
| *nme4* | -1.040 | 1.58E-02 |
| *rhag* | -0.869 | 1.59E-02 |
| *ctsba* | -0.610 | 1.59E-02 |
| *glsb* | -0.784 | 1.60E-02 |
| *vdra* | -0.863 | 1.60E-02 |
| *cited4a* | 1.041 | 1.63E-02 |
| *mpi* | 0.860 | 1.66E-02 |
| *asgrl1* | -0.634 | 1.74E-02 |
| *arhgap1* | 0.690 | 1.74E-02 |
| *hhla2a.1* | -1.482 | 1.74E-02 |
| *capn3a* | 1.355 | 1.76E-02 |
| *itga2.1* | -1.655 | 1.76E-02 |
| *lpcat3* | 0.766 | 1.77E-02 |
| *mfsd6a* | 1.071 | 1.80E-02 |
| *tbc1d10ab* | 0.865 | 1.85E-02 |
| *fstl3* | -1.567 | 1.85E-02 |
| *arhgap32a* | 0.970 | 1.85E-02 |
| *serpinb1l3* | 0.822 | 1.85E-02 |
| *pde1a* | 0.930 | 1.85E-02 |
| *slc25a25a* | 0.774 | 1.94E-02 |
| *igfbp5b* | -1.128 | 1.94E-02 |
| *aldh9a1a.1* | -0.906 | 1.94E-02 |
| *calcoco2* | -1.333 | 1.98E-02 |
| *slc7a9* | -1.194 | 2.02E-02 |
| *abcg2c* | -1.256 | 2.11E-02 |
| *atp1b1a* | 0.820 | 2.14E-02 |
| *ahsg2* | -1.228 | 2.16E-02 |
| *FAM53C* | 1.074 | 2.18E-02 |
| *lclat1* | 0.726 | 2.21E-02 |
| *ddit4* | 1.238 | 2.22E-02 |
| *scarb2c* | 0.730 | 2.23E-02 |
| *rfng* | 1.011 | 2.28E-02 |
| *ap2b1* | 0.588 | 2.29E-02 |
| *ELAPOR2* | 1.010 | 2.40E-02 |
| *tsc1b* | 0.936 | 2.40E-02 |
| *CR925731.1* | 1.310 | 2.40E-02 |
| *apoc4* | -1.103 | 2.41E-02 |
| *eef2a.1* | 0.885 | 2.55E-02 |
| *ncoa4* | 0.652 | 2.58E-02 |
| *si:ch211-196i2.1* | -1.591 | 2.60E-02 |
| *si:ch211-152c8.2* | 1.106 | 2.60E-02 |
| *nfil3-5* | -1.072 | 2.62E-02 |
| *dpp6b* | 1.494 | 2.67E-02 |
| *cfhl5* | -0.771 | 2.86E-02 |
| *cast* | -0.893 | 2.98E-02 |
| *gba* | 0.753 | 2.98E-02 |
| *cyp2k20* | -1.377 | 3.00E-02 |
| *ces3* | -1.088 | 3.04E-02 |
| *zgc:103759* | -1.436 | 3.05E-02 |
| *BX323800.1* | 1.324 | 3.05E-02 |
| *got2b* | 0.644 | 3.13E-02 |
| *nipsnap2* | 0.692 | 3.14E-02 |
| *st8sia6* | -0.970 | 3.19E-02 |
| *cyp2x9* | -1.206 | 3.20E-02 |
| *hgd* | -0.741 | 3.25E-02 |
| *plxna3* | 0.746 | 3.33E-02 |
| *plekhb2* | -1.198 | 3.45E-02 |
| *adipor1b* | 0.548 | 3.51E-02 |
| *cryaa* | 1.342 | 3.54E-02 |
| *igf2a* | 0.906 | 3.56E-02 |
| *nkiras2* | -0.985 | 3.56E-02 |
| *si:ch73-173p19.1* | -0.981 | 3.69E-02 |
| *pwwp2b* | 0.687 | 3.69E-02 |
| *CABZ01089907.1* | 1.364 | 3.72E-02 |
| *cds1* | 0.927 | 3.76E-02 |
| *pros1* | 0.934 | 3.76E-02 |
| *myrip* | -1.009 | 3.79E-02 |
| *ccdc32* | -0.957 | 3.79E-02 |
| *clptm1* | 0.857 | 3.88E-02 |
| *hmbsb* | -1.180 | 3.88E-02 |
| *FQ323156.1* | 1.232 | 3.93E-02 |
| *igf2b* | 1.131 | 3.97E-02 |
| *fosaa* | 1.383 | 3.97E-02 |
| *si:ch211-215a10.4* | 0.733 | 3.97E-02 |
| *lnx2b* | 0.646 | 3.97E-02 |
| *bach1a* | 0.911 | 3.98E-02 |
| *acadm* | -0.715 | 4.01E-02 |
| *slc26a3.2* | -0.642 | 4.02E-02 |
| *rgl1* | 0.677 | 4.02E-02 |
| *dio1* | 0.856 | 4.02E-02 |
| *tmem150aa* | 0.623 | 4.02E-02 |
| *CU207301.3* | 0.762 | 4.02E-02 |
| *anpepa* | -1.260 | 4.04E-02 |
| *herpud1* | 0.866 | 4.05E-02 |
| *loxl1* | -0.760 | 4.05E-02 |
| *chia.6* | -0.791 | 4.07E-02 |
| *lhpp* | 0.848 | 4.20E-02 |
| *cox8a* | 0.628 | 4.22E-02 |
| *sts* | 0.921 | 4.26E-02 |
| *xrcc1* | -0.815 | 4.27E-02 |
| *shroom4* | -1.051 | 4.27E-02 |
| *DOCK4* | 1.169 | 4.28E-02 |
| *si:ch211-120g10.1* | 1.367 | 4.37E-02 |
| *CR936540.1* | -0.963 | 4.37E-02 |
| *pip4k2aa* | 0.684 | 4.37E-02 |
| *prr5a* | -1.252 | 4.38E-02 |
| *pdxkb* | 0.966 | 4.39E-02 |
| *CR450764.11* | -1.244 | 4.51E-02 |
| *rnasel3* | -0.680 | 4.52E-02 |
| *spag6* | 1.333 | 4.52E-02 |
| *hikeshi* | 0.653 | 4.56E-02 |
| *CABZ01088308.1* | 1.132 | 4.62E-02 |
| *BX942813.1* | 0.947 | 4.69E-02 |
| *cyp27b1* | 1.209 | 4.72E-02 |
| *tnksa* | 0.594 | 4.76E-02 |
| *faah2a* | 0.812 | 4.81E-02 |
| *crispld2* | -1.362 | 4.83E-02 |
| *pdhb* | -0.549 | 4.86E-02 |
| *cyp46a1.3* | -1.268 | 4.87E-02 |
| *fam43a* | -0.829 | 4.90E-02 |
| *cicb* | -0.865 | 4.90E-02 |
| *ptpmt1* | -0.885 | 4.98E-02 |
| *cacna1db* | 1.330 | 4.98E-02 |
| *dhrs13l1* | -1.051 | 4.98E-02 |

**Supplementary Table 6d.** miR-21OE HCD v miR-21OE NCD

| **miR-21OE HCD v miR-21OE NCD** | | |
| --- | --- | --- |
| **Gene Name** | **L2FC** | **padj** |
| *hmgcra* | -3.668 | 7.86E-37 |
| *saa* | 0.126 | 5.13E-22 |
| *ASS1* | -3.008 | 9.75E-15 |
| *tmem59l* | 2.408 | 2.06E-08 |
| *dusp2* | 2.129 | 2.65E-07 |
| *chl1a* | 1.854 | 3.33E-07 |
| *cyp51* | -2.160 | 2.14E-06 |
| *si:cabz01040626.2* | 0.015 | 2.14E-06 |
| *egr2a* | 2.112 | 9.13E-06 |
| *si:ch211-284e136* | -1.991 | 2.27E-05 |
| *abcg1* | 1.894 | 5.22E-04 |
| *kcnh6b* | 2.163 | 5.94E-04 |
| *idh1* | -1.048 | 7.09E-04 |
| *abcg5* | 1.225 | 7.89E-04 |
| *CR450764.18* | -0.062 | 8.29E-04 |
| *kremen1* | 1.545 | 8.93E-04 |
| *rdh12* | -1.689 | 8.96E-04 |
| *relt* | 1.355 | 8.96E-04 |
| *zgc:86738* | -2.281 | 8.96E-04 |
| *crybb1l1* | -0.060 | 1.27E-03 |
| *fads2* | -1.381 | 1.28E-03 |
| *si:ch73-106k19.5* | -1.587 | 1.74E-03 |
| *dao.2* | -1.617 | 2.06E-03 |
| *mlxipl* | -1.028 | 2.82E-03 |
| *casp6b.2* | -0.064 | 3.30E-03 |
| *c4b* | 1.250 | 3.35E-03 |
| *si:ch211-28p3.4* | -0.097 | 3.35E-03 |
| *zgc:114181* | -0.190 | 5.15E-03 |
| *mb* | 1.459 | 6.36E-03 |
| *lss* | -1.158 | 1.12E-02 |
| *ppdpfa* | -1.362 | 1.47E-02 |
| *il1rapl1a* | 1.308 | 1.62E-02 |
| *CABZ01088367.1* | 1.225 | 1.62E-02 |
| *elovl2* | -1.202 | 1.66E-02 |
| *acss2l* | -1.346 | 1.73E-02 |
| *si:zfos-943e10.1* | -0.986 | 1.91E-02 |
| *sigmar1* | -1.102 | 2.32E-02 |
| *atic* | -0.962 | 2.32E-02 |
| *si:ch211-9d9.1* | 1.396 | 2.86E-02 |
| *zgc:66313* | -1.461 | 2.90E-02 |
| *msmo1* | -1.082 | 2.90E-02 |
| *spon2a* | 1.348 | 3.56E-02 |
| *ins* | -0.671 | 3.56E-02 |
| *selp* | 1.358 | 3.56E-02 |
| *NPC1L1* | -0.865 | 3.81E-02 |
| *Si:ch211-221f10.2* | 0.853 | 4.66E-02 |
| *nr4a1* | 1.211 | 4.83E-02 |
| *ptk6b* | -1.530 | 4.83E-02 |
| *ampd3a* | -1.338 | 4.96E-02 |
| *CR388178.1* | -0.732 | 4.96E-02 |
| *dusp5* | 1.398 | 4.97E-02 |
| *fosaa* | 1.446 | 4.97E-02 |
| *cox6a2* | 1.402 | 4.97E-02 |

*= GO cholesterol metabolism, cholesterol biosynthetic process, and/or cholesterol efflux

L2FC= log2 fold change

HCD= high cholesterol diet

NCD= normal control diet

**Supplemental Table 7:** Primers to amplify *dre-mir-21-1* with BbsI cut sites.

| **Primer Name** | **Sequence** |
| --- | --- |
| *dre-mir-21-1_BbsI_F* | *GTACGGGCTATGTCTTCccaccctctcctctcatcag* |
| *dre-mir-21-1_BbsI_R* | *GTACGGGCTATGTCTTCtggcgacgctaaaataagaca.* |

**Supplemental Table 8:** IDT gBlockTM Sequence with MfeI and BamHI Cut Sites for Creation of Dendra2 Sponge. Bolded sequences highlight enzyme cut sites and the 6 sponge sites are underlined.

| **Sponge name** | **Sequence** |
| --- | --- |
| *dre-miR-21SP* | *ctaATGAATG****CAATTG****TTGTTGTTgccaacaccctggataagctaGGAACCTTgccaacaccttggataagctaTCTGACGAgccaacacctatgataagctaCGCTTTAgccaacaccgtagataagctaGATCCTCGgccaacacctaggataagctaTATACCCGgccaacacctaagataagctaAACTTGTTTATTGCAGCTTATAATGGTTACAAATAAAGCAATAGCATCACAAATTTCACAAATAAAGCATTTTTTTCACTGCATTCTAGTTGTGGTTTGTCCAAACTCATCAATGTATCTTAAGGCGTAAATTGTAAGCGTTAATATTTTGTTAAAATTCGCGTTAAATTTTTGTTAAATCAGCTCATTTTTTAACCAATAGGCCGAAATCGGC****GGATCC****TGCGGCCatg* |

**Supplemental Table 9:** Primer sequences for qPCR

| **Gene** | **Primer name** | **Sequence** |
| --- | --- | --- |
| Lactate dehydrogenase | LDHD FWD  LDHD REV | GCCTGGGGTGACACGAAAAA  GGACTGCATTTGTGCCTGATG |
| Ferroptosis suppression protein 1 | FSP1 FWD  FSP1 REV | GGCTTTGACTTCATACAAAACAGAA  AAATGGAAGAGCGTGCAACC |
| Insulin-like growth factor (IGF) binding protein 2 | igfbp2a FWD  igfbp2a REV | TGTGACAAGAGGGGGCAGTA  AGGGGCTTAAAGGGCAAGAG |
| Inter-Alpha-Trypsin Inhibitor Heavy Chain 2 | Itih2 FWD  Itih2 REV | TGTGCAGATCCCCAAACGAG  CGTGCACTTCTGTCCGAAAC |
| vitamin D receptor | VDR FWD  VDR REV | GCTGATTAGCCTCGACTGTTCT  AATCCGGTGGCTTTGTCTCC |
| GABA Type A Receptor-Associated Protein | GABARAP FWD  GABARAP REV | AGGCGATCAGAGGGAGAGAA  CAGTCAGATCAGAAGGGACCAG |
| nicotinamide nucleotide transhydrogenase | nnt FWD  nnt REV | TTCCGGGTGACGGCTGTAAG  ACTAGTGAGGCGGTTGAGGG |
| Niemann-Pick disease type C2 protein | npc2 FWD  npc2 REV | CCACGCGTCCGAGGAAACT  AGTAAAACCACGCCGAGCA |
| Beta Actin | BACTIN FWD  BACTIN REV | CGAGCAGGAGATGGGAACC  CAACGGAAACGCTCATTGC |
