## Supplemental Figures for "microRNA-21 promotes dysregulated lipid metabolism and hepatocellular carcinoma"

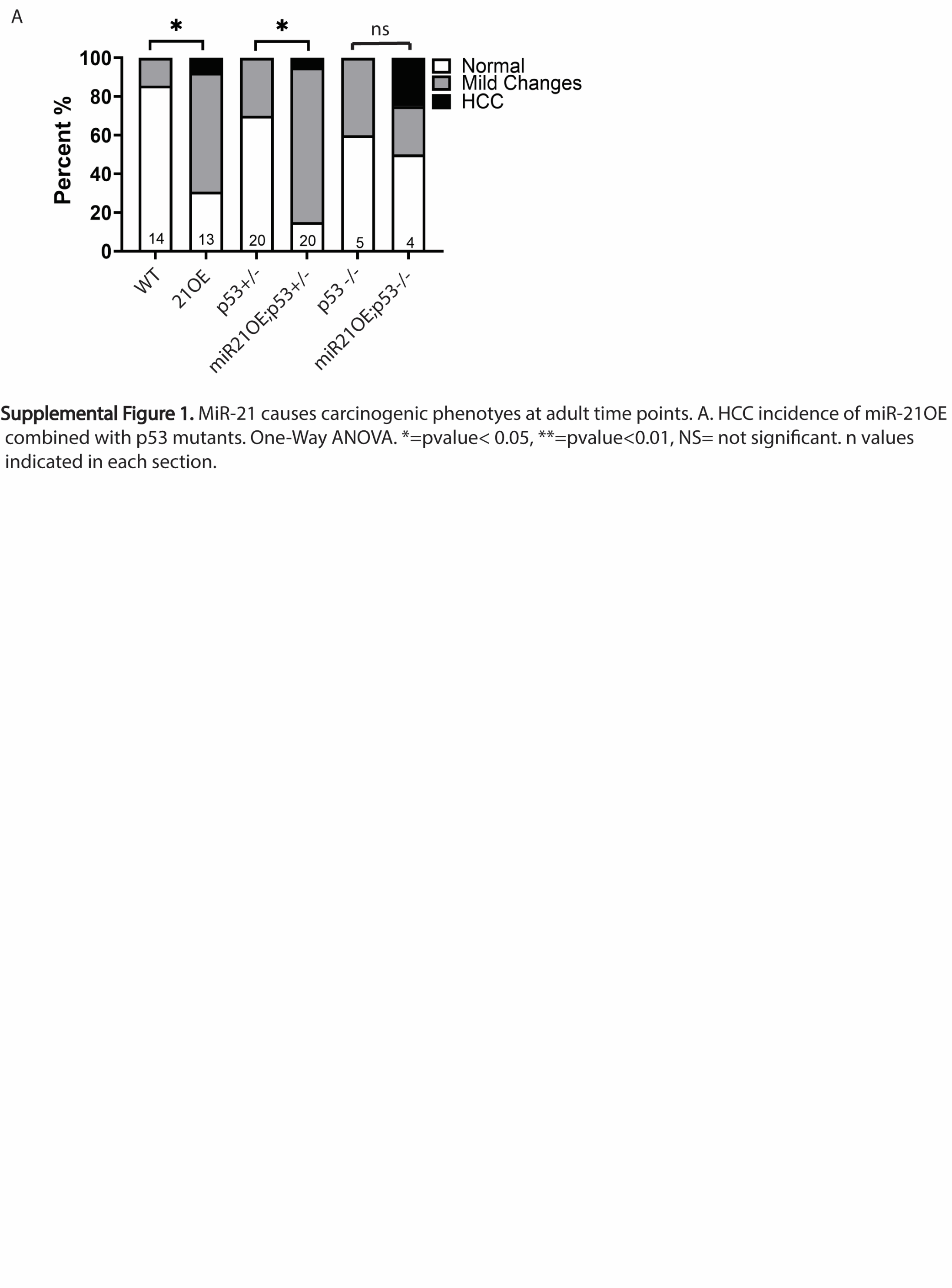

**Supplemental Figure 1. miR-21 causes mild changes and HCC in adult zebrafish.** Histology of miR-21OE (21OE) and non-transgenic wildtype sibling (WT) adult zebrafish livers (12 mpf) in the presence of p53 homozygous (p53-/-) or heterozygous (p53+/-) mutations was assessed by hematoxylin and eosin (H&E) stain. P values determined with GraphPad Prism, one-way ANOVA: ns, not significant; *, p < 0.05. N values are indicated at the bottom of each column.

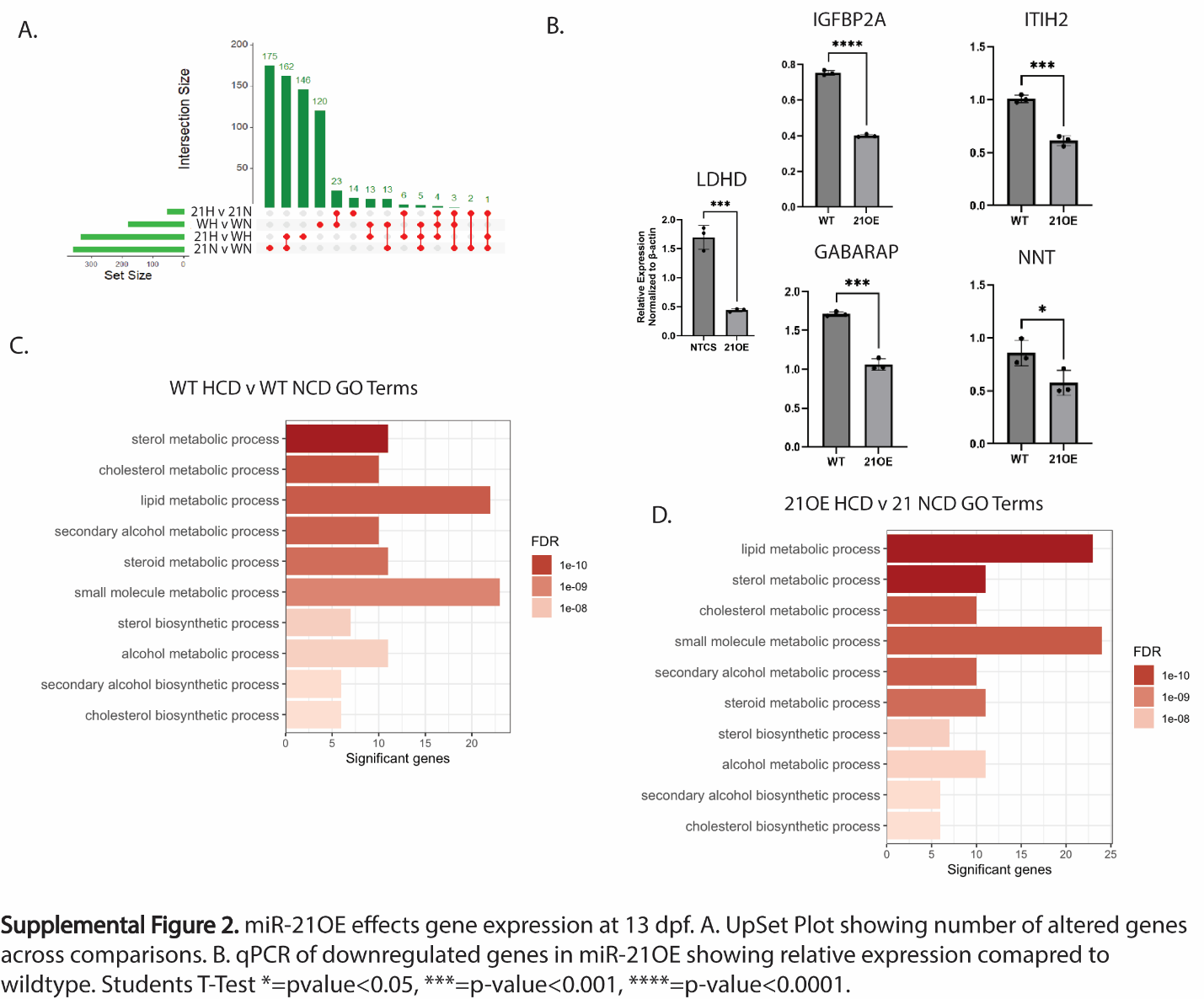
**Supplemental Figure 2. miR-21OE affects gene expression in larval zebrafish**. A. UpSet Plot of RNA-seq data showing number of altered genes across comparisons for miR-21OE (21) and non-transgenic wildtype sibling control (W) livers from 13-dpf zebrafish fed high cholesterol diet (H) or normal control diet (N). B. qPCR of genes in miR-21OE (21OE) and controls (W) relative to β-actin. Students T-Test *=pvalue<0.05, ***=p-value<0.001, ****=p-value<0.0001. C-D. Significant gene ontology (GO) terms for WT (C) and miR-21OE (D) on high cholesterol diet (HCD) compared to normal control diet (NCD), RNA-seq data.

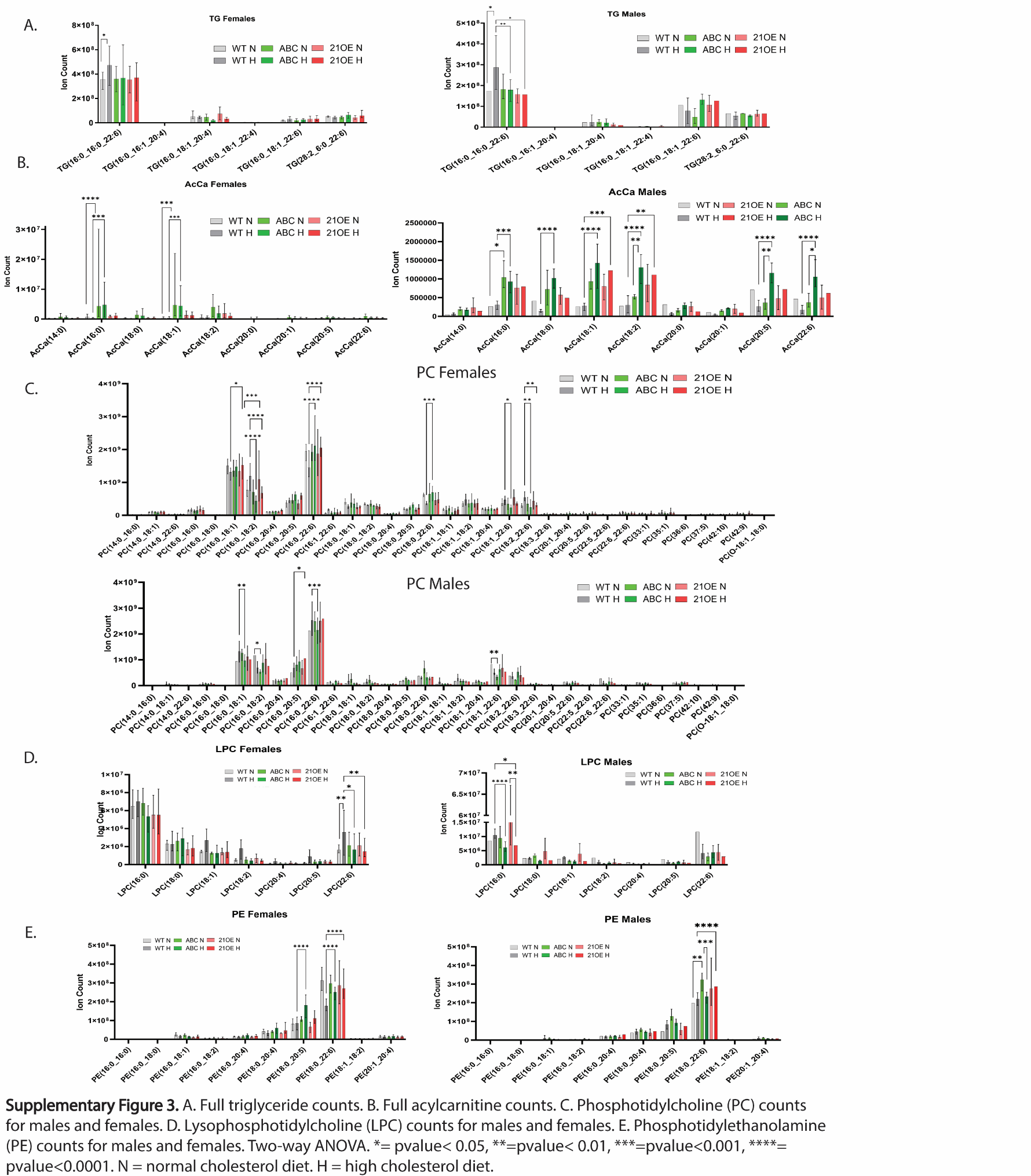
**Supplementary Figure 3. Lipidomics analysis of male and female adult zebrafish.** A. Triglyceride (TG) counts. B. Acylcarnitine (AcCa) counts. C. Phosphotidylcholine (PC) counts. D. Lysophosphotidylcholine (LPC) counts. E. Phosphotidylethanolamine (PE) counts. Two-way ANOVA: *, < 0.05; **, p < 0.01; ***, p < 0.001; ****, p < 0.0001. N = normal cholesterol diet. H = high cholesterol diet. WT = non-transgenic wildtype sibling control. ABC = activated β-catenin overexpression. 21OE = miR-21 overexpression.
